# p38 blockade reverses the immune suppressive tumor microenvironment in metastatic breast cancer

**DOI:** 10.1101/2025.05.08.652949

**Authors:** Priyanka Rajan, Robert Zollo, Yanqi Guo, Mohammed Alruwaili, Justin Zonneville, Mackenzie Lieberman, Brian Morreale, Caitlin James, Mark Long, Scott H. Olejniczak, Joseph Barbi, Scott I. Abrams, Andrei V. Bakin

## Abstract

Metastatic breast cancer (MBC) is a life-threatening disease with limited therapeutic options. The immune suppressive tumor microenvironment (TME) limits the potency of the antitumor immune response and facilitates disease progression and metastasis. Our current study demonstrates that p38α is a druggable target in the TME that regulates the outcome of the immune-tumor interaction. The study revealed that systemic blockade of p38α reduces metastasis, and this anti-metastatic response is negated by depletion of CD8^+^ T cells. Single-cell transcriptomic analysis of the immune-TME showed that pharmacological p38 inhibition (p38i) or tumor-specific inactivation of p38α by CRISPR/Cas9 (p38KO) resulted in a less exhausted and more activated CD8^+^ T cell phenotype. Immunophenotyping analyses demonstrated that p38 blockade reduced the expression of multiple inhibitory receptors on CD8^+^ T cells (i.e., PD-1, LAG-3, CTLA-4), indicating a reversal of immune exhaustion and enhanced immune activation systemically and in the TME. In contrast, p38 blockade did not exhibit inhibitory effects on T cells in proliferation assays *in vitro* and did not affect the proportion of regulatory T cells *in vivo*. The major negative impact of p38 blockade *in vivo* was on the myeloid populations, such as myeloid-derived suppressor cells (MDSCs) and tumor-associated macrophages (TAMs). Further, tumor p38α activity was required for the expression of cytokines/chemokines and tumor-derived exosomes with high chemotactic capacity for myeloid cells. Altogether, this study highlights a previously unrecognized p38α-driven pathway that promotes an immune suppressive TME and metastasis, and that therapeutic blockade of p38α has important implications for improving antitumor immunity and patient outcomes.

**STATEMENT OF SIGNIFICANCE:** This study highlights a previously unrecognized p38α-driven tumor pathway that promotes an immune suppressive microenvironment and metastasis, and that therapeutic blockade of p38α has important implications for improving antitumor immunity and patient outcomes.

## INTRODUCTION

Metastatic breast cancer (MBC) is a life-threatening disease with lowest 5-year survival rates in patients with metastatic triple-negative breast cancer (mTNBC) [1, 2]. A critical contributing factor to these poor survival outcomes in MBC patients is the hostile tumor microenvironment (TME) [3, 4], characterized by dense stroma and presence of immune suppressive myeloid populations that support cancer progression and resistance to multiple therapies [4, 5]. Thus, identifying druggable targets in the TME is important for improving treatment outcomes.

Tumor infiltrating myeloid-derived suppressor cells (MDSCs) and tumor associated macrophages (TAMs) facilitate tumor invasion and suppress anti-tumor immune responses leading to metastasis [6–8]. The molecular pathways that mediate tumor-induced recruitment and/or expansion of these myeloid cell populations represent potential therapeutic targets [4]. Accumulating evidence implicates p38 mitogen-activated protein kinase (p38 MAPK; p38) in tumor-induced changes in the TME [9–13]. Early studies have shown that the tumor p38-alpha isoform (p38α, *MAPK14*) contributes to the dissemination of TNBC by mediating tumor-intrinsic capacities that drive invasion and angiogenesis [9, 10, 12]. Recently, the pro-metastatic activity of p38α in mTNBC was linked to ability of these tumors to recruit CD163^+^ TAMs and to induce the expansion of MDSCs that facilitate pre-metastatic niches in the lungs and liver [11]. In align with this observation, p38α activity in the lung fibroblasts was also implicated in the recruitment of pro-tumorigenic neutrophils, supporting colonization of the lungs by melanoma cells [14]. Further, p38α activity in cancer-associated fibroblasts (CAFs) was also linked to changes in the metastatic TME and the anti-tumor immune response [15]. However, the mechanisms by which p38α supports expansion and recruitment of immune suppressive myeloid cells remain incompletely understood.

To that end, we investigated the contribution of p38α to altering the immune landscape of the TME using genetic inactivation of p38α (*Mapk14*) by CRISPR/Cas9 or pharmacological blockade in immune competent preclinical mTNBC models. The study provided new insights into the mechanism by which tumor p38 drove an immune suppressive TME and enhanced metastasis. Blocking p38 activity reduced metastasis and this response was CD8^+^ T cell dependent. Single-cell transcriptomic analysis of tumors revealed that p38 blockade led to a signature consistent with increased CD8^+^ T cell activation and reduced CD8^+^ T cell exhaustion. In addition, tumor-specific inactivation of p38α reduced the accumulation of immune suppressive myeloid cells in the TME, as well as in circulation. Mechanistically, p38α blockade reduced tumor-mediated chemotactic attraction of myeloid cells by reducing the levels of tumor-produced extra-cellular vesicles and chemokines that control the mobilization of such immune suppressive myeloid cells. Together, these findings highlight a previously unknown p38α-driven pathway that promotes a hostile TME, thereby, facilitating metastasis and identifies p38α as a potential therapeutic target in MBC.

## METHODS

### Cell lines and culture conditions

Human mammary carcinoma cell line MDA-MB-231 (RRID: CVCL_0062), mouse mammary carcinoma 4T1 cell line (RRID: CVCL_0125) and mouse monocyte-macrophage cell line RAW 264.7 (RRID: CVCL_0493) were obtained from American Type Culture Collection (ATCC, Manassas, VA, USA), and cultured as recommended by ATCC. 4T1 cell line was modified to express a Luciferase reporter (4T1-luc). MDA-MB-231-p38^AGF^ (dn-p38) cells were generated by retroviral transduction [9]. The cells were routinely screened for mycoplasma and all studies were made with mycoplasma-free cells. Cell cultures were maintained in media supplemented with 10% heat-inactivated fetal bovine serum (HI-FBS) and 1% penicillin/streptomycin at 37°C with 5-10% CO_2_ in a humidified incubator. All cell lines were authenticated using short tandem repeat profiling by ATCC or the Roswell Park Core within the last three years.

### p38 inhibitor, Antibodies and Other Reagents

The information is presented in Supplementary Information.

### CRISPR-Cas9 mediated p38 knockout (p38KO)

To inactivate p38α in 4T1-luc cells, two CrisprRNA targeting Exon 4 in mouse *Mapk14* (Transcript ID: ENSMUST00000114754.7) were designed using Crispor software [16], the sequences are provided in the Supplemental Information. CrisprRNA (crRNA) and trans-activating crRNA (tracrRNA) samples were mixed at 160μM each (1:1 ratio) and annealed by a touchdown PCR method as follows 95°C, 5min; chill to 55°C, 5C/min; to 25°C, 5 min. To form the functional ribonucleoprotein (RNP) complex, 61μM of recombinant Cas9-3NLS protein was added to the guide RNA complex and incubated at room temperature for 20 min. The RNP mixture was added to 3×10^6^ cells suspended in 100μl optiMEM media kept on ice, followed immediately by electroporation under optimized conditions in a NEPA21 electroporator (Nepa Gene Co. Ltd., Japan), and then transfer into 6-well plates containing 2 ml antibiotic-free complete medium. Single cell suspensions were subjected to single cell sorting of viable cells (7-AAD-negative) using a BD FACSAria II sorter (BD Biosciences) equipped with 488nm, 635nm, 405nm and 355nm lasers and controlled by FACS Diva 8.0.1 software. Cell clones were expanded, replica plates were made, and sequenced by using primers flanking the intended cut sites in the exon. Clones with ∼100% p38KO were selected for further analysis.

### Immunoblotting

Tumor cells were grown in a 6-well plate at 300,000 cells/well for 16 hours prior to treatment with 2μM p38i (ralimetinib, LY2228820), 10ng/ml TNFα or 2ng/ml TGFβ1 for indicated times. For UV irradiation, the cells were exposed to 100J/m^2^ or 250J/m^2^ using Stratalinker 2400 followed by 30 min incubation at 37°C, 5% CO_2_ before collection. Whole-cell lysates were prepared using NP40 lysis buffer containing PMSF, Na-Orthovanadate, and protease inhibitor cocktail. Protein concentration in lysates were quantitated and 30µg of protein/lane were resolved on SDS-PAGE gels. Proteins were transferred onto nitrocellulose membrane in 10% Methanol-SDS buffer and probed with appropriate antibodies. ECL reagent was used to visualize immune complexes on radiographic films.

### RT-qPCR

Cells were treated with 2µM p38i or vehicle DMSO for 24 hours. RNA was extracted using TRIzol reagent according to the manufacturer’s instructions. cDNA was prepared and qPCR was performed in triplicates as outlined in [10]. Primer sequences are present in Supplemental information.

### *In vivo* experiments

Female BALB/c mice (6-7-week-old) were purchased from the Charles River Laboratories (Wilmington, MA). Animals were kept at the Laboratory Animal shared Resource at Roswell Park in microinsular units and provided with food and water *ad libitum* according to protocols approved by the Institute Animal Care and Use Committee (IACUC). The facility is certified by the American Association for Accreditation of Laboratory Animal Care (AAALAC) and in accordance with current regulation and standards of the US Department of Agriculture and the US Department of Health and Human Services. For the CD8α depletion experiment, mice were implanted with 100,000 actively proliferating 4T1-luc or p38KO cells in the 4^th^ mammary fat pad in a 1:1 ratio of Matrigel and Dulbecco’s Phosphate Buffered Saline 1X (1X DPBS). Non-tumor bearing (naïve) mice were inoculated with 1X DPBS. Mice were monitored daily for signs of distress. Tumor volumes were measured using electronic calipers twice a week and calculated using the formula (length) x (width)^2^/2. Mouse weights were measured twice a week. On day 3 post implantation, mice were randomized into treatment groups based on mouse weights. Antibody to mouse CD8α subunit of CD8 and matched isotype control (IgG2b, κ) antibody were administered on day 3 by i.p injection at 200µg/mouse dose and weekly at 180µg/mouse thereafter. On day 4, treatments with vehicle (Veh, DPBS) or p38 inhibitor (p38i, ralimetinib, 30mg/kg) was initiated by daily oral gavage. On day 6 post-tumor inoculation, peripheral blood was collected by retro-orbital eye bleed in accordance with IACUC guidelines for flow cytometric verification of CD8^+^ T cell depletion. When the average tumor volumes reached 500mm^3^, mice were euthanized and tumor, lungs, liver, and spleen were collected.

For all the other experiments, mice were inoculated with 200,000 actively proliferating 4T1-luc control or 4T1-luc p38KO cells in the 4^th^ mammary fat pad in a 1:1 ratio of Matrigel and DPBS (n=6/group). On day 4 post-tumor inoculation, mice bearing control 4T1 tumors were randomized into vehicle and p38i treatment arms. The vehicle arm was administered with DPBS while the treatment arm received p38i (ralimetinib, 30mg/kg) by daily oral gavage. At day 14 post implantation, the mice were euthanized and subjected to necropsy and organ collection. Tissues were collected for RNA and protein analyses by snap-freezing in liquid nitrogen. Blood was collected for CBC by cardiac puncture.

### Complete Blood Count (CBC)

At the endpoint, blood was collected by cardiac puncture into EDTA solution to prevent coagulation. Analysis was performed using HemaTrue Analyzer and HeskaView Integrated Software (Version 2.5.2).

### Single cell RNA sequencing (scRNAseq)

Tumors from 3 mice per group were pooled together and minced in ice-cold 1X DPBS. The samples were enzymatically digested using a cocktail of active protease subtilisin A from *Bacillus licheniformis* and DNAse1 at 6°C for 30 min, to avoid collagenase and thermal stress induced transcriptional signatures [17, 18]. The samples were transferred onto gentleMACS C tubes and dissociated using the program ‘h_tumor_01’ on a gentleMACS dissociator [8]. The cell suspension was washed with ice cold 1X DPBS containing 10% FBS and then passed through 70μm and 40μm filters. The single cell suspension was resuspended in 1X DPBS containing 0.04% BSA and stained for CD45 and 7AAD. Live (7AAD-negative) CD45^+^ cells (∼200,000) were sorted from the suspension using BD FACS AriaII and resuspended in 1X DPBS containing 0.04% BSA at 10^6^ cells/ml. About 5,000 cells/group were subjected to scRNAseq procedure (Droplet-seq) using 3’ Rapid Amplification of cDNA Ends (3’ RACE) approach and 10X Genomics Chromium Controller v3.1 at the Roswell Park Genomic Shared Resource. Sequencing data were collected from 4,588 cells of the vehicle (Veh) group, 6,343 cells of the p38i group, and 5,212 cells of the p38KO group. A total of 32,285 genes from 16,143 cells were analyzed further by R programming to check for quality, identify immune cell types and changes in gene expression profiles.

### Flow cytometry

For the CD8^+^ T cell depletion study, cells obtained through retro-orbital eye bleed were treated with ACK lysis buffer at room temperature for 10 mins. The cells were washed with 1ml flow buffer and resuspended in 1ml flow buffer (0.1% BSA in 1X DPBS). Cells were stained with trypan blue and counted using hemocytometer. Approximately 500,000 cells in each sample were blocked with anti-CD16/32 and then stained with anti-CD45, anti-CD4, anti-CD8 and LIVE/DEAD fixable aqua at room temperature. The samples were acquired using Fortessa B.

For analysis of T cells, the Spleen and lymph nodes were crushed and filtered through a 70-μm nylon cell strainer (Corning). Red blood cells were lysed by ACK lysing buffer (Thermo Fisher Scientific). Tumors were cut into 2 to 3 mm pieces, then transferred into a gentle MACS C tube (Miltenyi Biotec). Enzymatic cocktail consisting of collagenase and hyaluronidase was added and the tissue was processed through a gentleMACS Dissociator (Miltenyi Biotec). Single-cell suspensions from tumors, spleens or tumor-draining lymph nodes (TdLNs) were obtained by passing dissociated tumors through 70-μm nylon cell strainers (Thermo Fisher Scientific). Cells were washed with flow running buffer (0.1% BSA in PBS, Thermo Scientific). The cells were stained with trypan blue and counted. About 1,000,000 cells were resuspended in staining buffer containing antibodies to CD16/CD32 (Fc receptor blocker, dilution, company) and T cell or myeloid cell surface markers (Supplemental Key Resources) after labelling non-viable cells using LD-Aqua (Molecular Probes, Eugene, OR, USA). After surface marker staining, cells were fixed/permeabilized using a FOXP3/transcription factor staining kit (eBiosciences) as per manufacturer’s protocol for intracellular staining. Cells were then stained with antibodies to T or myeloid cell internal markers (Supplemental Key Resources). For cytokine markers, the cells were stimulated with PMA, ionomycin and brefeldin A for 4 hours at 37 °C. This was followed by cell surface and intra-cellular staining as described above. All the flow data was collected on CytoFlex LX (Beckman Coulter) and analyzed using FlowJo v10.5.2 (RRID: SCR_008520).

### T cell stimulation

T cells were isolated from spleens of 6-week-old female BALB/c mice using Pan CD3+ T cell negative isolation kit. The cells were pre-treated with 0.5 and 2μM p38i (ralimetinib) for 1 hour, followed by TCR stimulation using αCD3/αCD28 coated plates using media containing 25 U/mL recombinant murine IL-2 (BioLegend). The cells were expanded for 3, 5 or 7 days in media containing 25 U/mL IL-2 in the presence/absence of p38i. Then, the cells were stained with anti-CD45, CD4, CD8, CD44 and CD62L. LD-aqua was used to exclude non-viable cells. To analyze the expression of IFNγ and TNFα, the cells were stimulated with PMA, Ionomycin and Brefeldin A along with p38i for 4 hours, followed by surface and intracellular staining.

### Luciferase assay

Mouse lung tissues were homogenized suspended in NP-40 lysis buffer containing protease inhibitor cocktail. Luminescence assays was done using Veritas Microplate Luminometer and Luciferase Assay System (Promega, Madison Wisconsin).

### Tumor cell conditioned media (TCM)

Tumor cells were plated on 100mm dishes and left to adhere overnight. The following day, when the cells were 80% confluent, the cells were washed 3 times with PBS and incubated in serum free media (SFM) supplemented with 2µM p38i (ralimetinib). After 48 hours, TCM was collected and centrifuged at 10,000xg for 20 mins. TCM was used immediately or stored at -80°C prior to use.

### Chemotaxis assay

RAW 264.7 cells (1×10^5^/well) were plated in SFM in the upper chamber of 8.0μm pore transwells (Corning) and were incubated with CM collected from tumor cells. After 4 hours, the cells were fixed in 100% methanol and cells remaining at the top of the membrane were removed using wet cotton swabs. Cells that had migrated through the pores to the lower surface were stained with 0.5% crystal violet. The membranes were washed and air-dried overnight. Four random images were recorded at 200x magnification using Nikon T100 microscope equipped with Spot 5.6 software and cells were counted.

### Chemotaxis assays with primary mouse T cells and tumor cell conditioned media (TCM)

Spleens were harvested from female 5-week-old BALB/c mice (two spleens per a biological replicate). CD3^+^ T cells were isolated, activated with anti-CD3/CD28-coated magnetic beads and cultured in the presence of IL-2 in DMEM media containing 10% FBS, 1% P/S, L-Glut, HEPES, and beta-mercaptoethanol (BME) for 4 days. The anti-CD3/CD28 beads were removed by magnetic separation. Cells were washed with PBS to remove serum and resuspended in serum-free DMEM. TCM media (600µl) from vehicle-control, p38i-treated, or p38KO 4T1 cells was added to the bottom chambers 15min prior cell seeding. T cells (∼0.5×10^6^) in 100µl serum-free DMEM were added to the top chamber of each transwell. After 8h, media from duplicate bottom chambers was collected and pooled, and then centrifuged at 1500rpm for 5 mins. The cell pellets were resuspended in 100ul SFM, and cells were counted using automated cell counter.

### scRNAseq data pre-processing and quality control

Cell Ranger package was used to align the reads to mouse reference genome and construct gene count matrices for each cell. All downstream analysis was performed in RStudio Desktop v1.4 equipped with R v4.2.1. The filtered feature matrix containing gene counts, cell barcodes and gene names was utilized to construct a SingleCellExperiment object using SingleCellExperiment package [19]. Per cell quality control was performed using the addPerCellQC function from Scater package [20]. Cells with sequencing depth less than 1500 total counts, less than 300 detected genes, and more than 20% mitochondrial content were removed. Next, Per Gene QC was performed by examining the mean vs. variance relationship of gene counts in the log_2_ counts/million scale. Genes with variance of less than 0.5 log_2_ counts per million were removed. Dimensionality reduction was performed to visualize the data points in a 2D plane using t-distributed Stochastic Neighborhood Embedding (t-SNE) method as outlined in Scater package [20]. Identity of immune cell types was determined using Single R package [21]. Mouse immunological gene expression data from Immunological Genome Project (ImmGen) database [22] was used as reference which was accessed using Celldex package [21]. Single R correlates the expression of cell type specific markers between reference and query data. About 20 different cell types (both immune and non-immune) were found to be present, from which non-immune cell types (Epithelial cells, Fibroblasts, Stem cells and Stromal cells) were excluded from analysis. In addition, the immune cells that were less than 1% abundant i.e. Basophils, B cells, pro-B cells, Mast cells and γδ T cells were excluded. The cell types that passed the criteria included T cells, Dendritic cells, Neutrophils, Monocytes, Macrophages, NK cells, NKT cells and Innate Lymphoid Cells. Seurat v5 was used for further analysis. In some of these cell populations, differential gene expression analysis was performed between the p38i and vehicle groups using FindMarkers function from Seurat using the following cutoffs: logFC > 0 and p value < 0.05. Heatmaps were constructed as outlined in ComplexHeatmap package [23] and in [24]. The pathway enrichment analysis was performed using the fgsea package [25] with NES defined as enrichment score normalized to mean enrichment of random samples of the same size.

### Statistics

Statistical analysis was performed using Graphpad Prism software (version 10.5) unless otherwise indicated. The data represent biological replicates (n) as mean value with standard error (mean +-SE). Statistical significance of data comparisons was determined using Students t-test with two tailed distribution, one-way ANOVA or two-way ANOVA with Tukey’s or Dunnett’s multiple comparisons test as indicated in figure legends. Statistical significance was achieved when p < 0.05.

## RESULTS

### CD8^+^ T cells contribute to the anti-metastatic activity of p38 blockade in MBC

Prior work showed that blockade of p38 reduces metastasis while increasing tumor infiltration by CD8^+^ T cells, although the role of CD8^+^ T cells in this response was not investigated [11]. Here, we examined the contribution of CD8^+^ T cells to the anti-metastatic effects of p38 blockade using a well-recognized mTNBC model of 4T1 mammary carcinoma cells in syngeneic BALB/c mice [26–29]. Female mice were implanted orthotopically with 4T1-Luc cells carrying a luciferase reporter and treated with a highly potent and selective inhibitor for α and β isoforms of p38 (p38i) Ralimetinib [30], or vehicle-control (**Fig. 1a**). At 100 mm^3^ tumor sizes, mice were subjected to administration of an anti-CD8 blocking antibody or a matched isotype IgG control. Flow cytometry analysis of the peripheral blood showed effective depletion of CD8^+^ T cells, while CD4^+^ T cells were unaffected (**Fig. 1b, Suppl. Fig. 1a-b**). In line with previous studies [11, 15], treatment with p38i reduced tumor growth compared to vehicle-control, whereas depletion of CD8^+^ T cells alleviated this anti-tumor effect of p38i (**Fig. 1c**, **Suppl. Fig. 1c**). The on-target activity of p38i was confirmed by assessment of phospho-MAPKAPK2 (*p*-MK2) levels in the tumor tissues (**Fig. 1d**). Evaluation of luciferase activity in the lung tissues showed that p38i reduced pulmonary metastasis, whereas depletion of CD8^+^ T cells negated this anti-metastatic effect (**Fig. 1e**). In a biological replicate, we observed that p38i reduced growth of the 4T1 tumors in the third week of treatment (**Suppl. Fig. 2a**) and significantly reduced metastasis (**Suppl. Fig. 2c**). Depletion of CD8^+^ T cells enhanced growth and metastasis of the 4T1 tumors treated with p38i or vehicle-control, and these effects were statistically significant (**Suppl. Fig. 2a-c**). Together, these findings demonstrated that CD8^+^ T cells are important effectors of the anti-tumor and anti-metastatic activities of systemic p38 blockade.

**Figure. 1.**
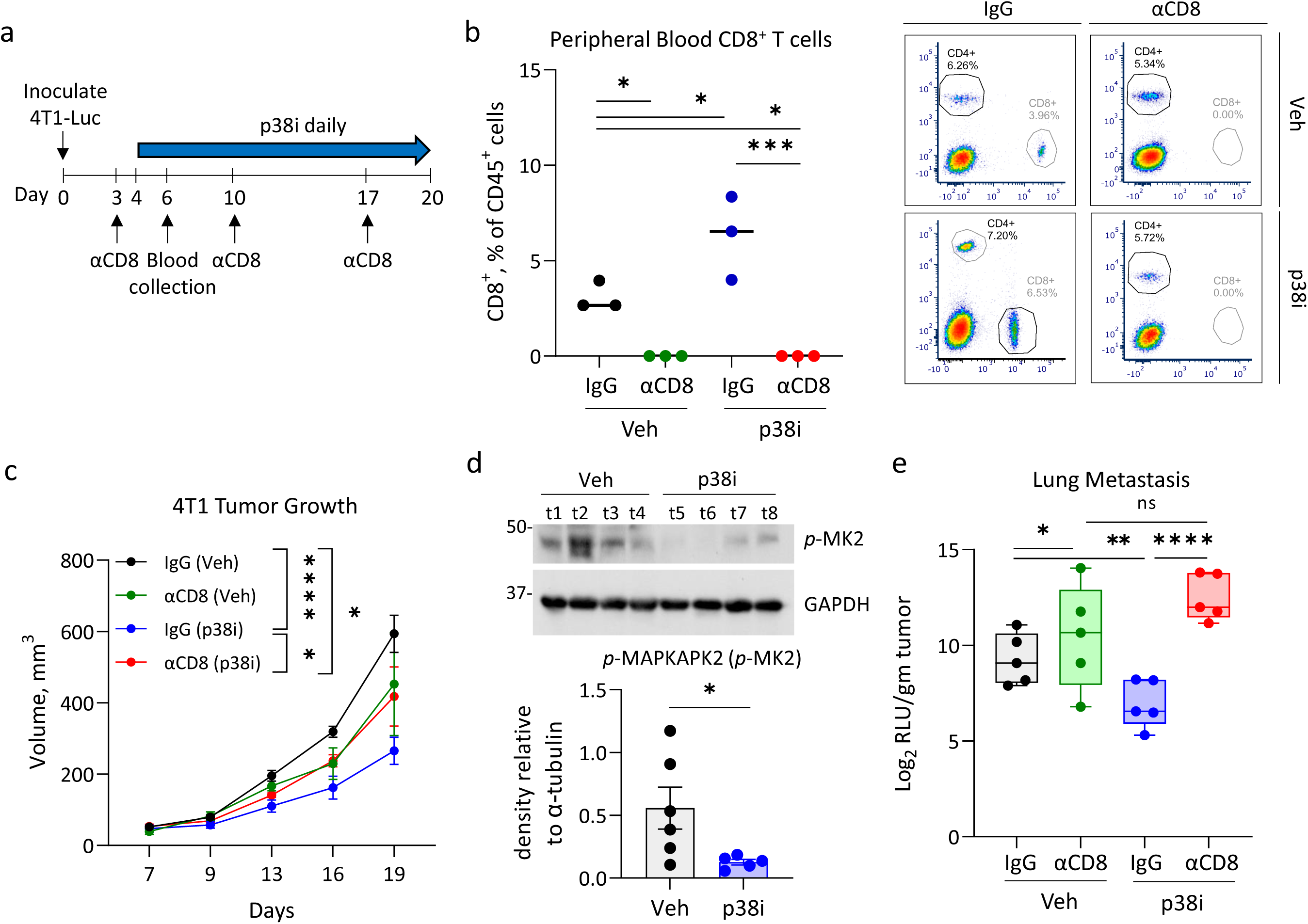
CD8^+^ T cells contribute to anti-metastatic activity of the systemic p38 blockade. (**a**) Experimental design of the *in vivo* study. Mouse mammary carcinoma 4T1-luc cells were implanted into mammary fat pads of BALB/c mice. On day 3, mice received i.p. injections of antibody to CD8. (**b**) Flow cytometric evaluation of CD8^+^ T cell levels in peripheral blood of mice (n=3/group) along with representative flow plots. Plots show %CD4^+^ and %CD8^+^ of live CD45^+^ cells. Comparisons were performed using one-way ANOVA. (**c**) Tumor growth curves for the individual groups. Error bars indicate Mean +/- SEM; the group comparisons were done using two-way ANOVA. Statistical values are shown at day 19 post implantation. (**d**) Immunoblots of whole tumor extracts from mice bearing 4T1 tumors from the vehicle control and p38i treated groups. Statistical analysis was done by t-test. (**e**) Luminescence activity in whole cell extracts from the lungs of tumor bearing mice, n=6/group. Values are log_2_ transformed and normalized to average tumor weight. RLU-Relative luminescence Units. The CD8-depletion experiment was repeated two times. Comparisons were performed using t-test. *, p<0.05; **, p<0.01; ***, p<0.001; ****, p<0.0001; ns, non-significant.

### p38 blockade and T cell proliferation and differentiation

Next, we investigated whether p38i exhibit a direct effect on T cells using *in vitro* T cell activation assays [31]. Splenic CD3^+^ T cells were isolated from naïve BALB/c mice and stimulated with αCD3/αCD28 antibodies in the presence of vehicle-control or p38i (**Suppl. Fig. 3a**), using doses that inhibit p38-dependent responses in tumor cells (**Suppl. Fig. 3b-c**). After 3, 5, or 8 days in culture, the T cells were treated with PMA and ionomycin in the presence of brefeldin A to stimulate cytokine production. The cells were immune-stained and analyzed by flow cytometry. The analysis showed only a limited effect of p38i on the levels of CD8^+^ and CD4^+^ T cells (**Suppl. Fig. 3d**, gating strategy in **Suppl. Fig. 3j**). To assess the naïve and memory T cell status, the cells were stained for activation marker CD44 and lymphoid tissue homing receptor CD62L [32, 33]. Flow cytometry analysis showed an increased frequency of CD44^high^ CD62L^+^ CD8^+^ cells upon p38i treatment (**Suppl. Fig. 3e-g**), indicating a shift towards a central-memory like phenotype. This observation aligns with a reduction in levels of CD8^+^ T cells expressing IFN-γ or TNF at days 5 and 8 (**Suppl. Fig. 3h-i**). Together these data indicate that p38i does not affect proliferation of T cells while exhibiting a mild effect on a central memory like population of CD8^+^ T cells.

### Inactivation of p38 MAPK alters the immune TME

To better define the contribution of tumor-specific p38α activity to systemic p38i effects in the immune TME, the p38α isoform (encoded by *Mapk14*) was inactivated in 4T1 cells using the CRISPR/Cas9 approach (**Suppl. Fig. 4a**). Disruption of *Mapk14* was validated by sequencing and three knockout (p38KO) clones were selected for further testing. In all three p38KO clones, p38α protein was not produced, while other p38 isoforms were not affected (**Suppl. Fig. 4b**). Phosphorylation levels of p38α downstream targets CREB1 and ATF1 were reduced in the p38KO clones whereas the levels of phospho-ERK1/2 were not affected (**Suppl. Fig. 4c**). p38α isoform is known to mediate signaling in response to TGF-β1 and UV [9, 34]. Depletion of p38α reduced phosphorylation of CREB1 and ATF1 induced by TGF-β1 and UV compared to control cells (**Suppl. Fig. 4d-e**). Conversely, the phosphorylation levels of SMADs and ERKs were not affected in p38KO clones (**Suppl. Fig. 4d-e**), in agreement with the fact that these signaling events in response to TGF-β1 and UV are not mediated by p38α [9, 34]. p38KO clones and control cells (untreated or p38i-treated) showed comparable proliferation in cell culture (**Suppl. Fig. 4f**). Next, 4T1 and 4T1-p38KO cells were implanted into BALB/c mice to examine the tumor growth. To explore the effect of CD8^+^ T cells, mice were also treated with antibodies to CD8 or a matched isotype IgG control as described in Fig.1. The growth of the p38KO tumors was markedly reduced compared to the 4T1 tumors, while depletion of CD8^+^ cells increased tumor growth but the effect was marginal (**Suppl. Fig. 4g-h**). Depletion of CD8^+^ T cells (αCD8) accelerated growth of the 4T1 tumors compared to IgG-control (IgG). The spleen weights in the p38KO tumors were significantly lower than in the 4T1 tumors (**Suppl. Fig. 4i**). The growth defect in p38KO tumors is consistent with a strong anti-tumor effect of dominant-negative p38^AGF^ mutant (dn-p38) in the breast cancer MDA-MB-231 model observed *in vivo* but not in cell culture [10]. Together, these results further indicated that the anti-tumor effects of p38 blockade are related to tumor-host interactions *in vivo*.

To assess the impact of the p38 blockade on the immune TME, female BALB/c mice were implanted with 4T1 p38WT or p38KO cells. Mice with palpable tumors in the p38WT cohort (day 4) were divided in two groups and treated with vehicle or p38i for 10 days (**Suppl. Fig. 5a**). At day 14 post implantation, tumors from 3 mice per group were pooled together and single-cell suspensions were prepared and stained for CD45. Live CD45^+^ cells (negative for 7AAD) were sorted by flow cytometry and subjected to scRNA-seq (**Suppl. Fig. 5a**). The cell type populations were defined using Immunological Genome database [22] and SingleR package in R [21]. This cell type analysis identified the presence of T cells, Granulocytes (Neutrophils), Monocytes, Macrophages, Natural Killer (NK cells), Natural Killer T cells (NKT), Dendritic cells (DC) and Innate Lymphoid cells (ILC) in the CD45^+^ populations of the TME (**Fig. 2a**). Granulocytes represented over 50% of CD45^+^ immune populations in the tumor cohorts (**Fig. 2a**), in line with a robust granulocytic response in the 4T1 tumor model [26, 35, 36].

**Figure. 2.**
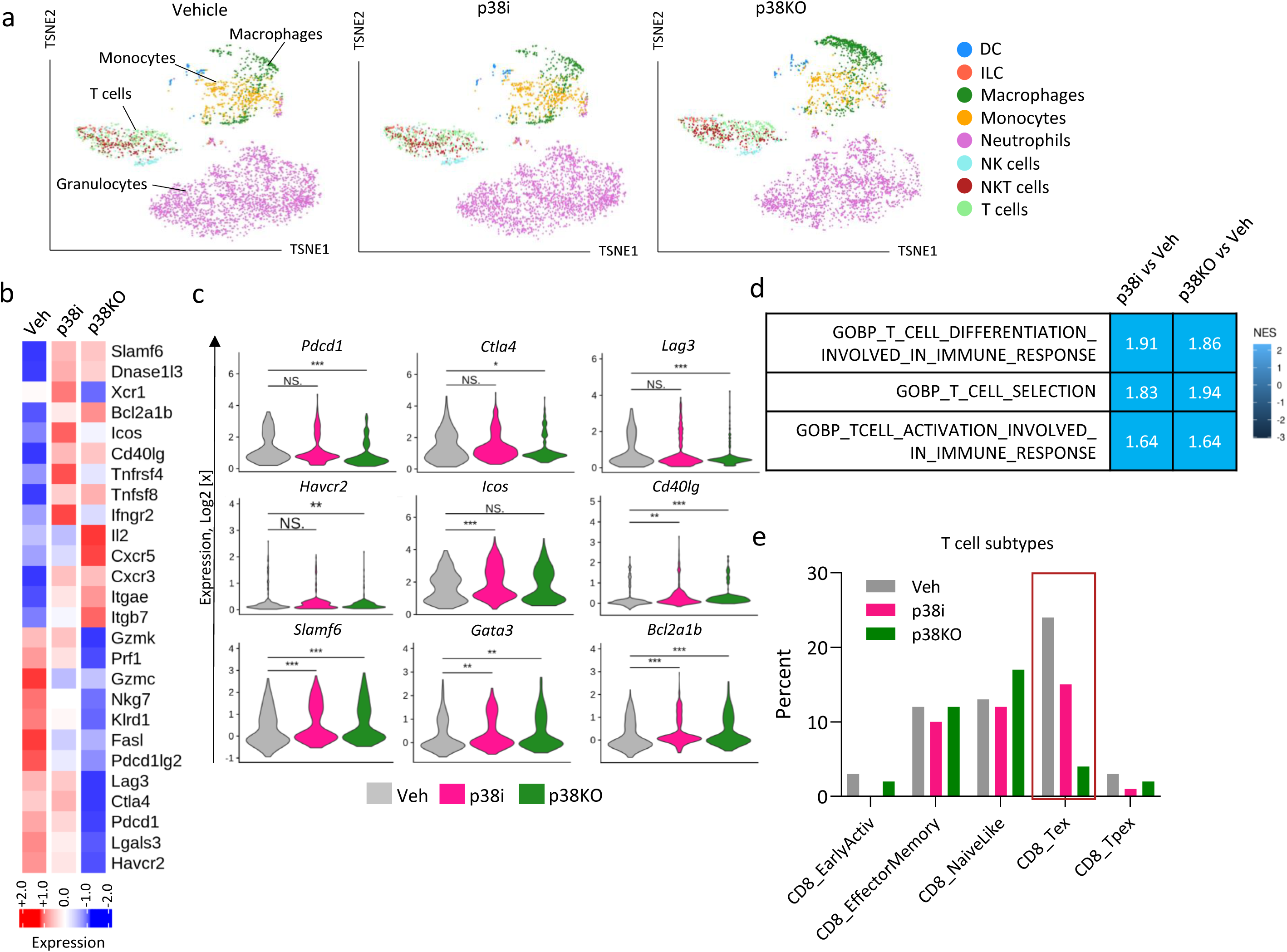
Inactivation of p38 alters the tumor immune microenvironment. The analysis of the tumor immune landscape was performed using single cell RNA sequencing (scRNAseq) in female BALB/c mice implanted with 4T1-luc (control and p38KO) cells. At day 4, mice bearing 4T1-luc tumors were treated with vehicle or p38 inhibitor (30mg/kg ralimetinib) for 10 days. At day 14 post implantation, the tumors were harvested and live CD45^+^ cells were subjected to scRNAseq. (**a**) The t-SNE plots show the immune populations identified in the tumors. (**b**) The heatmap of differential gene expression in T cells from the control and p38 blockade groups. Cutoffs used: p<0.05 and log2FC>0.5. (**c**) Violin plots for differentially expressed genes. Comparisons were performed using t-test. (**d**) Pathway enrichment analysis for T cells comparing the vehicle group to the p38i or p38KO groups; NES, normalized enrichment score. (**e**) CD8^+^ T cell subtypes detected by ProjecTILs package in R. *, p<0.05; **, p<0.01; ***, p<0.001; ***, p<0.001; NS, non-significant).

Transcriptomic analysis of tumor-infiltrating leukocytes showed that systemic p38 blockade with p38i and tumor-specific p38α inactivation decreased expression of exhaustion-associated transcripts (i.e., *Lag3, Ctla4, Pdcd1, Lgals3*, *Pdcd1lg2*, *and Havcr2*) in T cell populations, defined by expression of the T cell markers (**Fig. 2b-c**). The analysis of the exhaustion markers (e.g., Lag3, Ctla4, Pdcd1, Havcr2) in T cell subsets further confirmed reduction in mRNA levels of exhaustion markers in CD8^+^ and CD4^+^ T cell subsets in tumors from the p38 blockade groups (**Suppl. Fig. 5**). Consistent with these observations, the subtype analysis of T cells, using the ProjecTILs R package [37], showed a notable decrease in the exhausted T cell populations (CD8 T_ex_) in the p38i and p38KO groups compared to the vehicle-control (**Fig. 2e**). The pathway analysis showed the enrichment in the transcripts of ‘T cell differentiation involved in immune response’ and ‘T cell selection’ pathways in the p38i and p38KO groups (**Fig. 2d**). Further, p38 blockade increased T cell populations with high expression of genes associated with T cell activation and survival such as *Bcl2a1b, Ifn*γ*r2, Icos, Cd40lg,* and *Tnfrsf4* (**Fig. 2b-c**). In addition, the transcriptomic analysis showed upregulation of T cell transcription factors involved in differentiation to T-helper (Th) cells (i.e., *Slamf6, Gata3,* and *Ror*α) (**Fig. 2b-c**). The T cell subtype analysis showed an increase in CD4^+^ T cell levels, particularly the T-helper-1 populations in the p38i and p38KO groups compared to vehicle control (**Suppl. Fig. 5b-c**). Together, these findings revealed that blockade of p38 (by tumor-specific p38α inactivation or with p38i) generates a T cell molecular signature consistent with an activating and less exhausted immune phenotype.

### Immunophenotyping of T cell populations in tissues from tumor-bearing mice

To validate the transcriptomic data, the T cell populations infiltrating the primary tumor were assessed at day14 post-implantation using flow cytometry (**Fig. 3a-h**) with a gating strategy shown in **Suppl. Fig. 6a**. The analysis showed a reduction in CD8^+^ T cell populations expressing CTLA4 or LAG3 inhibitory receptors in the p38 blockade (p38i and p38KO) groups compared to the vehicle control (**Fig. 3a-b**), indicating a decrease in exhausted CD8^+^ T cell populations.

**Figure. 3.**
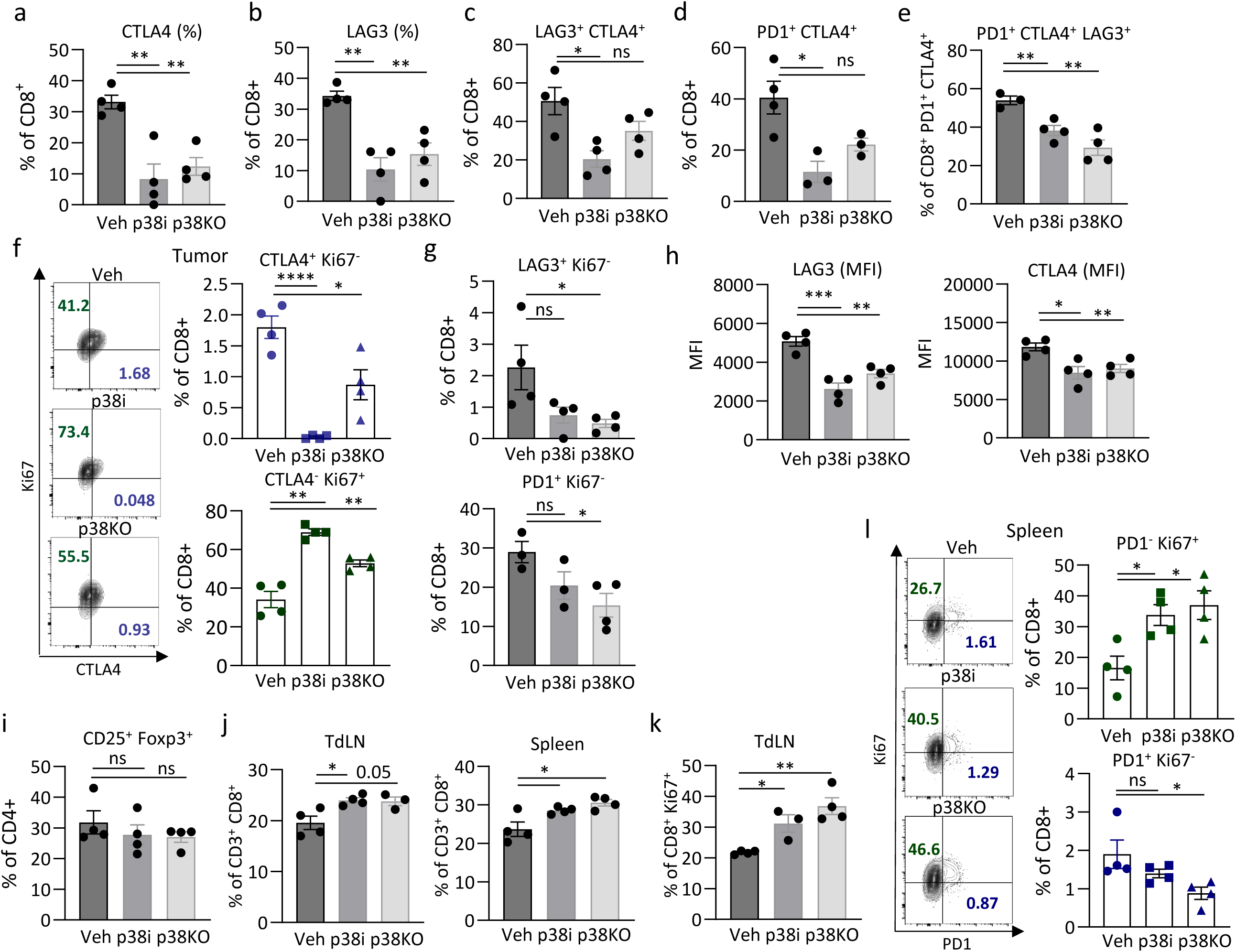
Immunophenotyping of T cells in the tumor and distant immune compartments. 4T1-luc control or p38KO cells were implanted into female BALB/c mice. At day 4, mice bearing control 4T1-luc tumors were treated with vehicle (Veh) or p38i (30mg/kg ralimetinib) by daily gavage. At day 14 post implantation, the tumors, lymph nodes, and spleens were harvested and subjected to the flow cytometry analysis. The **a-i** panels show the flow cytometry data for the tumor groups. (**a-b**) Percentage of CD8^+^ T cells expressing CTLA4 or LAG3. (**c**) Percentage of LAG3-positive CD8^+^ T cells co-expressing CTLA4. (**d**) Percentage of PD1-positive CD8^+^ T cells co-expressing CTLA4. (**e**) Percentage of LAG3-positive CD8^+^ T cells co-expressing CTLA4 and PD1. (**f**) Scatter plots and percentage graphs for CD8^+^ T cells expressing CTLA4 and Ki67 in the tumors. (**g**) Percentage of LAG3^+^ Ki67^-^ or PD1^+^ Ki67^-^ in the CD8^+^ T cell populations in the tumors. (**h**) Expression levels of LAG3 or CTLA4 (in mean fluorescence units, MFI) in CD8^+^ T cells. (**i**) Percentage of T regulatory cells (CD4^+^ CD25^+^ Foxp3^+^) in the tumors. (**j**) Percentage of CD8^+^ T cells in the TdLNs and spleens. (**k**) Ki67-positive (%) CD8^+^ T cells in the TdLNs. (**l**) Scatter plots and percentage graphs for CD8^+^ T cells in the spleens that Ki67-positive and PD1-negative (PD1^-^ Ki67^+^, proliferating and non-exhausted) or Ki67-negative and PD1-positive (PD1^+^ Ki67^-^, exhausted and non-proliferation). Representative data of 3 biological repeats are presented as mean +/- SEM for n=3-4 mice/group. Comparisons were performed by t-test (*, p<0.05; **, p<0.01; ***, p<0.001; ****, p<0.0001; NS, no significance).

Furthermore, the proportions of CD8^+^ T cells expressing multiple inhibitory receptors (CTLA4, LAG3, and PD1) were also decreased in the p38i and p38KO tumors compared to the control (**Fig. 3c-e, Suppl. Fig. 6b**). This observation is consistent with a notion that exhausted CD8^+^ T cells may express multiple inhibitory receptors that act *via* various inhibitory pathways [38]. Conversely, the proportions of CD8^+^ T cells positive for proliferation marker Ki67 and negative for CTLA4, PD1 or LAG3 were increased in the p38i and KO groups (**Fig. 3f-g, Suppl. Fig. 6a**), indicating a shift towards a more proliferative phenotype in CD8^+^ T cells. The absolute number and percentage of CD8^+^ T cells were also increased by p38 blockade (**Suppl. Fig. 6c-d)**. The effects on LAG3 and CTLA4 were also observed at the mean fluorescence intensity (MFI) of the inhibitory receptors, indicating a reduction in their expression levels (**Fig. 3h**). Oppositely, the p38 blockade did not affect the proportion of tumor infiltrating regulatory T cell (Treg) populations expressing CD4^+^CD25^+^Foxp3^+^ markers (**Fig. 3i**), suggesting that the observed effects on CD8^+^ T cell exhaustion are not related to Treg populations. Thus, these findings demonstrate that tumor-specific p38α inactivation or systemic p38i blockade resulted in a reversal of exhaustion in tumor-infiltrating CD8^+^ T cells, in agreement with the transcriptomic scRNAseq data.

Next, we examined T cell populations in the tumor draining lymph nodes (TdLNs) and spleens (**Fig. 3j-l, Suppl. Fig. 7-8**, and gating strategy in **Suppl. Fig. 7a-b**). Flow cytometry analysis showed increased proportions of CD8^+^ T cells in CD3^+^ leukocytes in the spleens and TdLNs from the p38i/p38KO groups compared to the vehicle control (**Fig. 3j)**, while levels of CD3^+^ leukocytes were also higher (**Suppl. Fig. 7c**). Furthermore, the proportions of Ki67^+^ CD8^+^ T cells were markedly increased in the p38 blockade groups in the TdLNs (**Fig. 3k, Suppl. Fig. 7d**) and spleens (**Suppl. Fig. 7f**), indicating enhanced proliferation of CD8^+^ T cells. In the spleens, a marked increase in proliferating PD1-negative CD8^+^ T cells (PD1^-^ Ki67^+^) were observed in the p38 blockade groups compared to the vehicle-control (**Fig. 3l**), while minor changes were observed for PD1^+^ Ki67^+^ CD8^+^ T cells accounting for only a small fraction of CD8^+^ T cells (**Suppl. Fig. 7e**). These findings indicate that the p38 blockade shifts T cells from an exhausted to more proliferative and activated state at distant lymphoid tissues, in addition to the primary TME.

To assess whether the p38 blockade affects memory T cells, CD44 and CD62L (L-selectin) were evaluated in CD8^+^ T cells in the TdLNs and spleens. These markers define major subsets of memory T cells: naive (CD44^low^CD62L^high^; T_naive_), central memory (CD44^high^CD62L^high^; T_CM_), and effector/memory (CD44^high^CD62L^low^; T_EM_) [32]. The analysis showed an increased proportion of CD8^+^ T cells expressing surface receptor CD62L^+^ in the TdLNs and spleens from the p38 blockade groups (**Suppl. Fig. 8a**). A higher frequency of CD44^high^ CD62L^+^ T_CM_-like cells and CD44^low^ CD62L^+^ T_naive_-like cells were also observed in the TdLNs and spleens from the p38 blockade groups compared to the vehicle-control (**Suppl. Fig.8b-c**). Furthermore, the absolute number of T_naive_-like and T_CM_-like cell populations were also increased in the TdLNs and spleens from the p38 blockade groups compared to the vehicle-control (**Suppl. Fig.8d-e**). These data indicated that blockade of p38 directed T cells in the peripheral lymphoid tissues towards phenotypes resembling naïve and central memory-like states, rather than egress of effector populations from the lymphoid tissue to the tumor site. Memory-like cells possess self-renewing properties and are linked to improved anti-tumor immunity [39]. These observed effects of p38 blockade may contribute to the enhanced anti-tumor CD8^+^ activity (**Figs. 2, 3**) and underlie the anti-metastatic p38i action **(Figs. 1c, e)**.

### Changes in myeloid populations caused by p38 blockade in tumor-bearing mice

Next, we investigated how the p38 blockade impacts myeloid cell populations identified in the scRNAseq study (**Fig. 2**). Transcriptomic analysis of monocytes showed that the p38 blockade decreased mRNA levels of markers associated with pro-tumorigenic monocytic and macrophage populations, *i.e.*, *Clec4e, Pla2g7, Cxcl2, Vegfa, Tgfb1, Ptgs2* and *Cd274* (**Fig. 4a-b and Suppl. Fig. 9a-b**). Notably, the expression of the chemokine receptors (*Cxcr1, Ccrl2, Cxcr4, Ccr5*) and the receptors for monocyte-stimulatory cytokines (*i.e*., *Csf1r, Csf2ra,* and *Csfr2b*) were also reduced by p38 blockade (**Fig. 4a-b and Suppl. Fig. 9a-b**). Furthermore, p38i enhanced macrophages expressing *Sphk1, Il7r, Tnfrsf9* which is indicative of a shift towards an anti-tumor phenotype (**Suppl. Fig. 9a-b**).

**Figure. 4:**
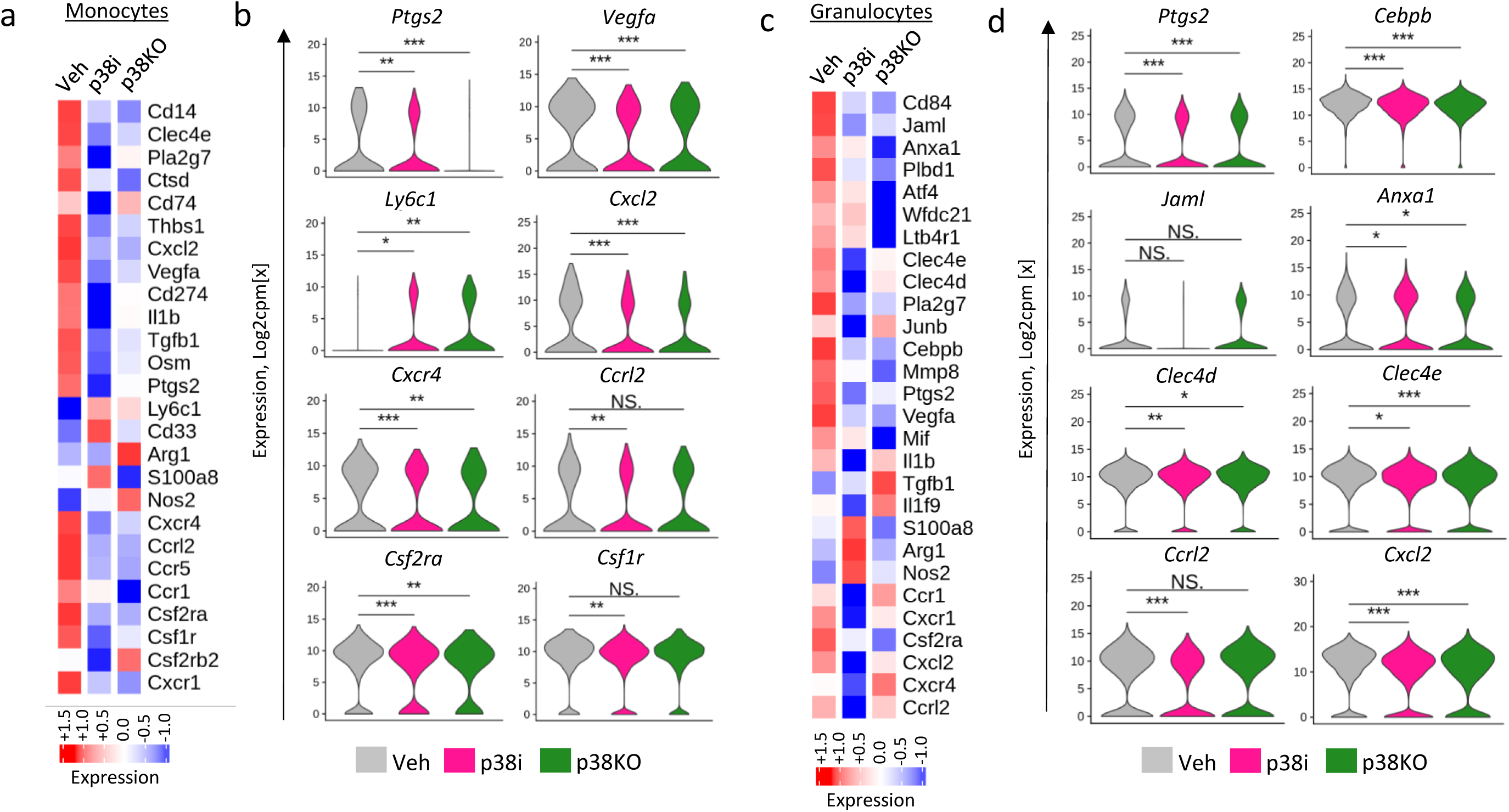
Effects of the p38 blockade on the transcriptome of monocytes and granulocytes. (**a**) The heatmap of differential gene expression in monocytic cell populations (scored in log2 counts per million) from the scRNAseq analysis using the vehicle, p38i-treated, and p38KO tumor groups. (**b**) Violin plots show distribution of gene expression levels, scored in log2 counts per million, in the monocytes from the vehicle, p38i, and p38KO tumor groups. Comparisons by t-test. (**c**) Heatmaps show scaled average gene expression levels for granulocytic MDSC populations (scored in log2 counts per million) from the vehicle, p38i, and p38KO tumor groups. (**d**) Violin plots show distribution of gene expression levels (in log2 counts per million) in the granulocytes from the vehicle, p38i, and p38KO tumor groups. Comparisons using t-test: *, p<0.05; **, p<0.01; ***, p<0.001; ****, p<0.0001; NS, non-significant).

Analysis of granulocytic populations showed a marked reduction in the expression of markers of PMN-MDSCs (i*.e.*, *Cd84, Jaml, Wfdc21, Ptgs2, Cebpb, Vegfa*) (**Fig. 4c-d**). Cd84 and *Jaml* were recently identified as surface receptors on MDSCs, with CD84 expressed by both human and mouse MDSCs [40]. *Ptgs2* (encoding COX2) is involved in the expansion and function of MDSCs [6, 41], while the transcription factor *Cebpb* is important for the generation of MDSCs [42]. In addition, blockade of p38 diminished expression of the matrix metalloprotease *Mmp8* in granulocytes (**Fig. 4c**). Mmp8 produced by PMN-MDSCs enhances metastasis by promoting extravasation and engraftment of circulating tumor cells [43]. Systemic p38 blockade with p38i decreased expression of the chemokine receptors *Ccrl2, Cxcr1, Csf2ra, and Cxcr4* in granulocytes, while only marginal changes were observed in the p38KO group (**Fig. 4c-d**). This observation suggests a direct effect of p38i on granulocytes that reduces their chemotactic capacity. Taken together, the transcriptomic data indicate that systemic p38 blockade with p38i or tumor-specific inactivation of p38α reduced markers contributing to expansion of monocytic and granulocytic populations with immune-suppressive characteristics.

To gain further insights, myeloid cell populations were examined in the spleen of tumor-bearing mice, a known site of extramedullary hematopoiesis (EMH) in tumor-bearing mice [11, 44, 45] and in patients with advanced-stage solid cancers [46]. The spleens were markedly enlarged in 4T1 tumor-bearing mice of the vehicle-control group compared to naïve mice (**Fig. 5a**). Pharmacological blockade of p38 reduced spleen sizes, in agreement with a prior report [11]. Furthermore, tumor-specific inactivation of p38α also reduced spleen sizes to values observed in the naïve group (**Fig. 5a**), indicating that tumor-derived signals regulate enlargement of the spleens. Analyses of the peripheral blood showed that blockade of p38 (by p38i and p38KO) diminished the levels of monocytes and granulocytes in the circulation (**Fig. 5 b, c**). The assessment of markers associated with immune-suppressive activities in splenic granulocytes (CD11b^+^Ly6G^+^) showed elevated mRNA levels of *Il10, Vegfa, Tgfb1,* and *Arg1* in the tumor-bearing vehicle-control group compared to naïve mice (**Fig. 5d-g**). In contrast, the expression levels of these markers were markedly reduced in the p38-blockade groups, approaching levels seen in the naïve mice group (**Fig. 5d-g**). Notably, low mRNA levels of *Il12a* were observed in splenocytes from the tumor-bearing mice, whereas splenocytes from the naïve and p38-blockade groups exhibited high mRNA levels of *Il12a* (**Fig. 5h**). IL-12a is an alpha subunit of IL-12, IL-17, and IL-23 heterodimeric cytokines that are implicated in regulation cell-based immunity including T cell activation, differentiation, effector functions, and memory [47].

**Figure 5.**
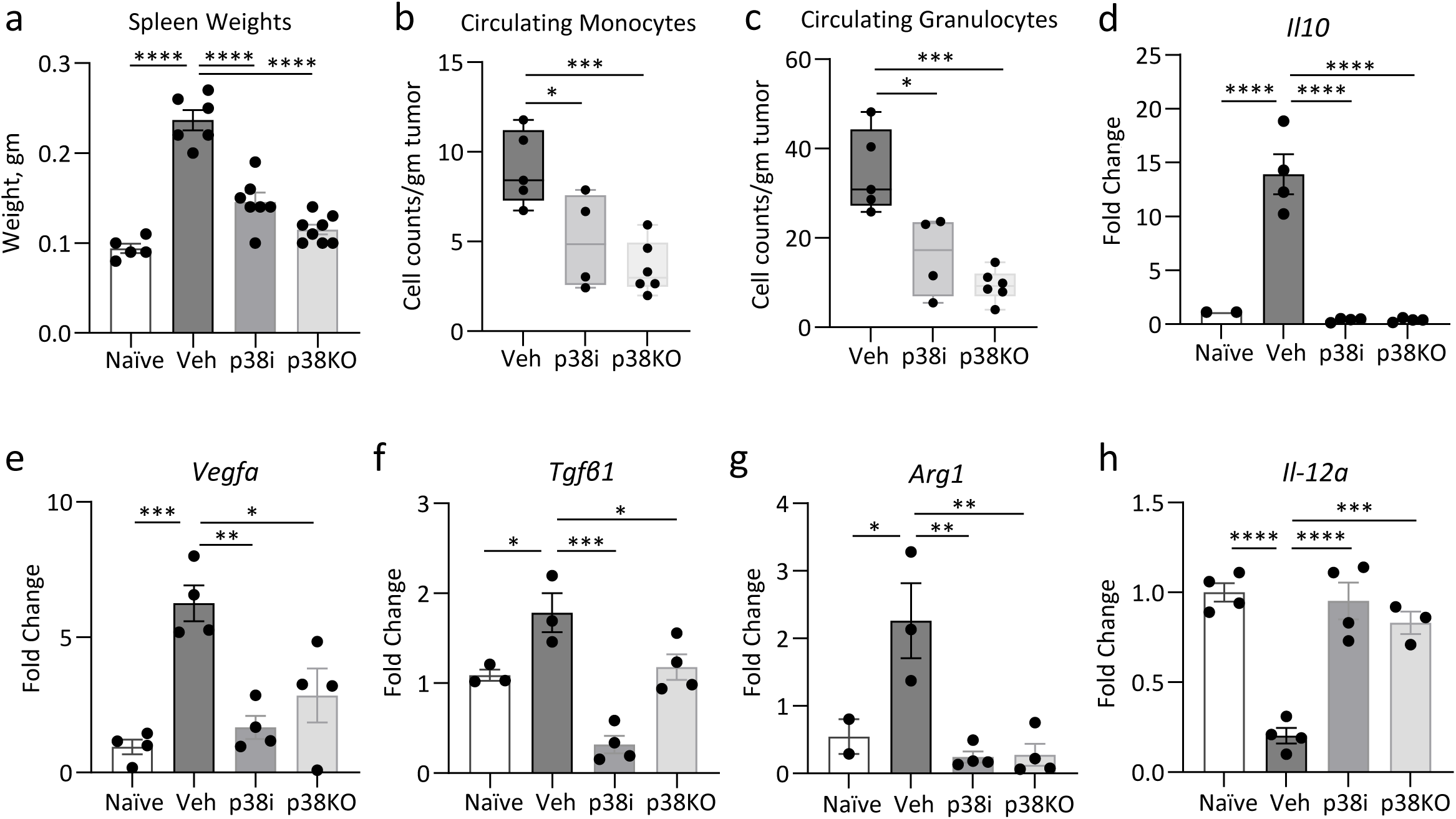
Effect of the p38 blockade on myeloid cell populations in the spleen. Tumor 4T1-luc control or 4T1-p38KO cells were implanted into mammary fat pads of BALB/c mice. At day 4, mice bearing 4T1-luc tumors were treated with vehicle or p38i (20mg/kg ralimetinib) by daily oral gavage. At day 14 post implantation, the spleens were harvested and Gr1^+^ splenocytes were isolated and subjected to RT-qPCR. (**a**) Weights of spleens from the vehicle, p38i, and p38KO groups. (**b-c**) Levels of monocytes and granulocytes in the peripheral blood are shown relative to tumor size. Comparisons were done by t-test. (**d-h**) mRNA levels of target genes in Gr1^+^ splenocytes. The fold changes relative to the vehicle group are shown as mean +/- SEM for 3 independent experiments. Comparisons by one-way ANOVA: *, p<0.05; **, p<0.01; ***, p<0.001; ****, p<0.0001.

To further validate these results, the flow cytometry analysis was performed for spleens and tumors. Analysis of myeloid populations in spleens showed that the p38 blockade reduced splenic granulocytes (CD11b^+^Ly6G^+^) expressing markers of immunosuppressive activities (Arg1, iNOS, VegfA) at both the proportion and intensity levels (**Suppl. Fig.10, gating strategy Suppl. Fig.12**). Further, p38 blockade increased the frequency of CD101+ granulocytes, indicating a shift towards a mature phenotype characterized by CD101 expression [48]. Moreover, the flow cytometry analysis of tumors showed that the p38 blockade (p38i or p38KO) reduced inflitration of CD11b^+^ myeloid cells and granulocytic Ly6G^+^ Ly6C^lo^ populations in the tumor (**Suppl. Fig.11**, gating strategies for the spleen and tumor are shown in **Suppl. Fig.11**). Likewise, p38 blockade increased the frequency of CD101^+^ mature granulocytes. These data demonstrate that blockade of p38 exhibits two effects characterized by a shift towards a less immunosuppressive phenotype and a reduction in numbers of suppressive myeloid cells.

Together, these findings indicate that systemic p38 blockade and tumor-specific inactivation of p38α markedly reduce the generation and/or expansion of pro-tumor myeloid cell populations, while positively regulating factors promoting T cell mediated immunity.

### Tumor p38 controls production of chemotactic cytokines and chemokines

Thus far, our results suggest that tumor p38α activity regulates factors that stimulate expansion and recruitment of pro-tumor myeloid cells (TAMs, MDSCs) by affecting myelopoiesis in the spleen. Prior studies showed that inhibition of p38 in splenocytes did not interfere with the generation of MDSCs in cell culture settings [11]. Thus, p38 may exhibit pro-myeloid action indirectly *via* chemotactic activity of secreted cytokines and chemokines. This hypothesis was examined by testing the chemotactic capacity of tumor cell conditioned media (TCM) using transwell chemotaxis assays with the mouse RAW264.7 macrophage cell line. The TCM samples were prepared by incubating tumor cells in serum-free media for 48 hours. TCM from 4T1 cells strongly induced transwell migration of myeloid cells compared to the serum-free media controls (**Fig. 6a** and **Suppl. Fig. 13a**). In contrast, transwell migration was markedly diminished in TCM from the p38i and p38KO groups (**Fig. 6a** and **Suppl. Fig. 13a**). Comparable results were obtained with TCM from the human TNBC cell line MDA-MB-231 expressing EGFP or a dominant-negative (dn) p38^AGF^ mutant (**Fig. 6b** and **Suppl. Fig. 13b**). To address a potential impact of p38i or non-peptide small molecules (e.g., prostaglandins), chemotaxis assays were performed with TCM depleted for compounds smaller than 3,000Da using Amicon filters. The chemotaxis assays showed that filtered-TCM stimulated migration of myeloid cells while this response was reduced in the p38 blockade groups (p38i and dnp38, **Suppl. Fig.13c-d**). These results supported a role of chemotactic cytokines in the chemotactic activity of tumor cells. Further, chemotactic activity of TCM towards T cells was measured in primary mouse T cells activated by anti-CD3/anti-CD28 treatment (**Suppl. Fig. 14a**). It was observed that the migration of activated T cells was enhanced by TCM from p38i-treated or p38KO cells compared to TCM from vehicle-treated 4T1 cells. This opposite effect of p38 blockade on TCM-mediated migration suggest that p38 may control expression of tumor secreted factors, which can negatively or positively regulate the chemotactic migration of T cells.

**Figure 6.**
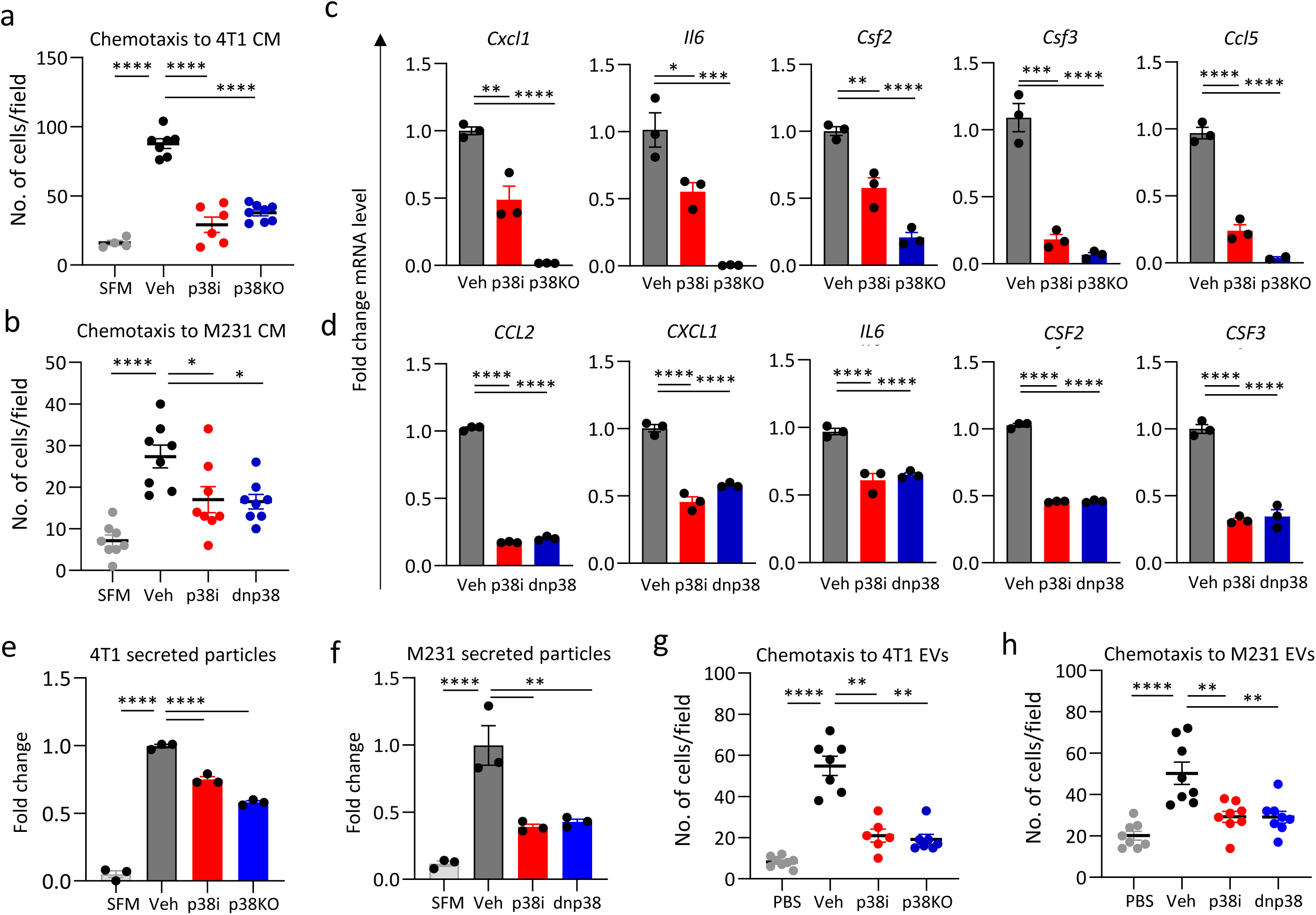
The contribution of the tumor p38 signaling to the tumor-myeloid crosstalk. Tumor cell conditioned media (TCM) was prepared using mouse mammary tumor 4T1 cells or human breast adenocarcinoma MDA-MB-231 cells incubated for 48hrs in serum-free nedia with or without p38i. Exosomes were isolated from TCM using an ultracentrifugation approach. Chemotactic capacities of TCMs and exosomes were evaluated in transwell migration assays using mouse macrophage cell line RAW 264.7. (**a**) Chemotactic capacity of TCMs from 4T1 cells in the vehicle, p38i and p38KO groups. (**b**) Chemotactic capacity of TCMs prepared from MDA-MB-231 cells treated with vehicle (Veh, DMSO) or p38i (2µM ralimetinib), or MDA-MB-231 cells expressing dn-p38 (AGF) construct. Serum-free media served as a negative control. (**c**) mRNA levels of cytokines and chemokines expressed by 4T1 cells (treated with vehicle or p38i for 24h) or p38KO cells. Fold changes relative to vehicle-treated cells using 5S rRNA for normalization. (**d**) mRNA levels of cytokines/chemokines in MDA-MB-231 cells: vehicle, p38i-treated (2µM ralimetinib) for 24hrs, or dn-p38. Fold changes relative to vehicle-treated cells using GAPDH for normalization. (**e**) Extracellular vesicles from 4T1 TCMs were analyzed using Nanoparticle Tracking Analyzer. Fold differences show the number of the secreted particles relative to the vehicle-treated cells. (**f**) Concentration of EVs in TCMs prepared using MDA-MB-231 cells (see description panel **b**). Fold differences show the number of the secreted particles relative to the vehicle-treated cells. (**g-h**) Chemotactic capacity of EVs prepared from the 4T1 cells (**g**) or MDA-MB-231 cells (**h**) treated with vehicle (Veh, DMSO) or p38i (2µM ralimetinib), or MDA-MB-231 cells expressing dn-p38 (AGF) construct. Serum-free media served as a negative control. The representative data are shown from 3-6 independent repeats. Data are shown as mean -/+ SEM. Comparisons by one-way ANOVA: *, p<0.05; **, p<0.01; ***, p<0.001; ****, p<0.0001.

Next, we examined whether p38 controls the expression of chemotactic factors. RT-qPCR analyses of 4T1 cells (**Fig. 6c**) showed that p38 blockade reduced mRNA levels of multiple cytokines and chemokines implicated in chemotaxis of myeloid cells. This observation was further validated in MDA-MB-231 cells (**Fig. 6d**). We also examined whether p38 blockade alters the expression of chemokine receptors in T cells infiltrating 4T1 tumors using our scRNAseq data. The analysis showed that mRNA levels of chemokine receptors were not altered in the p38i or p38KO tumor groups compared to the control (**Suppl. Fig. 14b**).

Recent evidence implicates extracellular vesicles produced by tumor cells, such as exosomes, as an intercellular form of communication to expand and mobilize immune-suppressive myeloid cells [49–52]. We then assessed whether blockade of p38 regulates EV production by tumor cells. Nanoparticle analysis of TCM showed that the sizes of EVs from mouse and human tumor cells range from 50-200 nm (**Suppl. Fig. 13e-f**). Inactivation of p38 (by p38i, p38KO, or dn-p38) did not change the sizes of the particles secreted by mouse and human tumor cells (**Suppl. Fig. 13e-f**) but markedly diminished the amounts of tumor-secreted particles (**Fig. 6e-f** and **Suppl. Fig. 13g-h**). Next, we examined the chemotactic capacity of EVs secreted by the tumor cells. Transwell migration assays showed that the addition of 10^6-8^ EVs is sufficient to stimulate chemotaxis of mouse myeloid cells (**Suppl. Fig. 13i**). Then, the chemotactic activity of EVs from mouse and human tumor cells was assessed using 10^6^ particles from control cells and cells with inactivated p38 (p38i, p38KO, or dn-p38). Chemotactic capacity of EVs from p38-inactivated cells was markedly lower compared to vehicle-treated control cells (**Fig. 6g-h, Suppl. Fig. 13j, k**). Thus, p38 blockade affected both secretion of EVs and the chemotactic capacity of EVs. The latter finding is likely associated with changes in the EV content in alignment with a reduced expression of chemokines and cytokines (**Fig. 6c-d**).

Next, we explored the clinical relevance of p38-related markers in human breast cancer. The analysis of TCGA data showed an elevated expression of p38-related markers in the basal-like subset of breast cancer (i.e., TNBCs) (**Fig. 7a**). Kaplan-Meier survival curves showed a clinically significant difference in overall survival in TNBC patients with high expression levels of p38-related markers (**Fig. 7b**). The median survival for patients with basal-like breast tumors in the low expression group is 196.36 months *vs* 84.73 months in the high expression group. These differences were limited to TNBC patients as evaluation of patients with Luminal A, Luminal B or Her2^+^ breast cancer subtypes based on expression of p38-related markers did not show clinically significant difference (**Suppl. Fig. 15**). Together, these results provide evidence in favor of the model that tumor p38α activity promotes expression of cytokines and chemokines and secretion of chemotactic EVs that stimulate expansion of myeloid cell populations and their recruitment to the tumor and lungs (**Fig. 7c**). These myeloid populations facilitate metastasis in part by exhibiting immune suppressive activities on cytotoxic T cells, while blockade of p38α reduces T cell exhaustion and enhances T cell activation locally and systemically.

**Figure 7.**
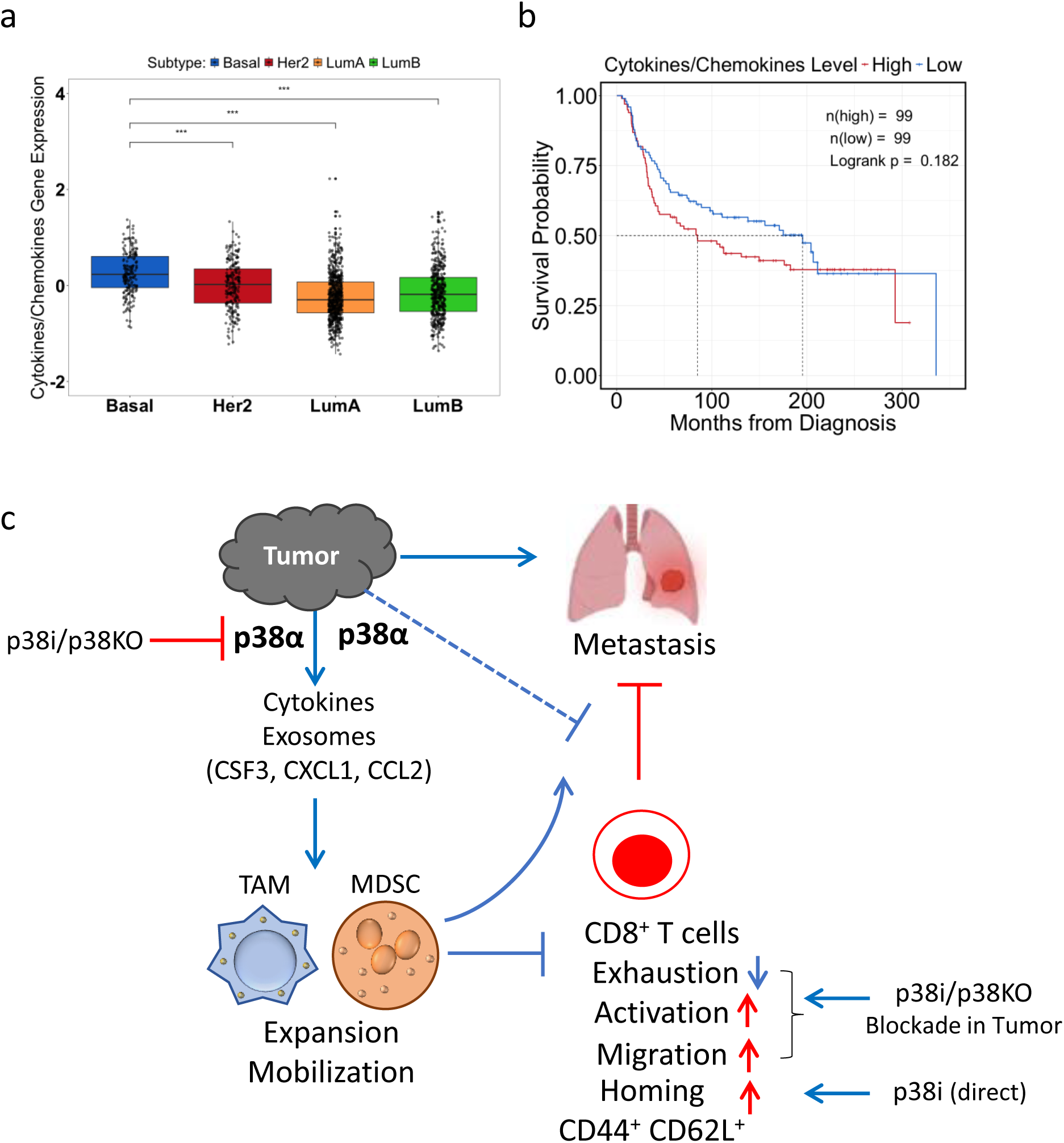
Expression of p38-regulated markers predicts survival in human breast cancer subtypes. (**a**) mRNA expression levels of cytokine/chemokine factors regulated by p38α in breast cancer subtypes. (**b**) Kaplan-Meier survival plots in patients with basal-like breast cancer using the TCGA database and stratified for high (red line) or low (blue line) expressions of cytokines and chemokines regulated by p38α. Median survival in the low expression group is 196.36 months vs 84.73 months in the high expression group. (**c**) A working model of tumor p38α contribution to tumor progression and metastasis by facilitating the production of tumor-derived factors which promote the expansion and mobilization of immune-suppressive TAMs and MDSCs. These myeloid cells are recruited to primary tumor and distant metastatic sites, where they facilitate formation of immune-suppressive microenvironment and inhibit anti-tumor immune response. In addition, systemic p38 blockade by p38i may exhibit a direct effect on CD8^+^ T cells enhancing their homing potential.

## DISCUSSION

The immune suppressive TME limits the potency of the antitumor immune response, facilitating disease progression and metastasis [3, 4]. Therefore, discovering druggable targets in the TME to reverse immune suppression is critical for improving patient outcomes. While prior research has implicated p38α in regulating the outcome of the immune-tumor interaction, the mechanistic details of how this occurs remained poorly defined. Here, we demonstrated that systemic blockade of p38 activity reduced metastasis, and this response was strongly dependent upon CD8^+^ T cells, as depletion of CD8^+^ T cells negated this anti-metastatic action (**Fig. 7c**). The impact of p38 blockade on reshaping the immune-TME landscape was further interrogated by scRNA-seq using either pharmacological (p38i) or tumor-targeted inactivation of p38α (p38KO). Comprehensive analyses revealed that both p38 blockade strategies led to an enhanced activation and less exhausted CD8^+^ T cell phenotype. These findings were validated at the protein level by immunophenotyping, including the analyses of the systemic T cell response within the tumor-draining lymph nodes (TdLNs) and spleens of tumor-bearing mice. Moreover, p38 blockade (*via* p38i or p38KO approaches) elicited a major impact on the myeloid compartment, reducing TAMs and MDSCs. Further investigation showed that tumor p38 regulates the expression of cytokines/chemokines and the production of exosome-like vesicles exhibiting chemotactic capacity. Blockade of p38 in tumor cells reduced their chemotactic activity towards myeloid cells, while enhancing migration of activated T cells. Together, our work provides new insights into the biology of tumor p38α and its actions on the immune-tumor interaction. Our findings unveiled a novel crosstalk among tumor cells, myeloid cells, and CD8^+^ T cells, regulating antitumor and anti-metastatic activity. This study thus extends our knowledge of how tumor p38α contributes to immune-tumor outcome.

Among the four isoforms of p38, the p38α isoform (encoded by *Mapk14*) is ubiquitously expressed at high levels [34]. Prior research showed that tumor p38α contributes to the invasive and metastatic capacities of tumor cells [9, 53], and tumor angiogenesis [10]. Importantly, a recent study revealed that tumor p38α plays a critical role in tumor-induced expansion of immune suppressive myeloid cells (TAMs, MDSCs), while depletion of MDSCs enhances tumor infiltration by cytotoxic CD8^+^ T cells [11]. Suppressive myeloid cells, MDSCs and TAMs, have been implicated in the induction of an exhausted T cell phenotype in cancer [54, 55]. MDSCs can suppress proliferation of T cells [56] and promote tumor angiogenesis and metastasis in mouse models [57]. The current study provides insights into the mechanisms by which tumor p38α contributes to immune suppressive environment and metastasis.

It was observed that the anti-tumor effects of p38 blockade by p38i or p38KO are dependent on the CD8^+^ T cell response, and depletion of CD8^+^ T cells negates this tumor control. The transcriptomic analysis and the immunophenotype studies in tumors showed that p38 blockade reduced the expression of multiple inhibitory receptors characterizing the T cell exhaustion phenotype (e.g., Ctla4, Lag3, Pdcd1, Havcr2). Blockade of p38 reduced the levels of exhausted T cells and increased levels of activated T cells within the tumor and spleen (as a systemic site). The levels of tumor-infiltrating CD4^+^CD25^+^Foxp3^+^ Treg cells were not affected by p38 blockade. Exhaustion of T cells is among major limiting factors of immunotherapy efficacy [58], and reactivation of T cells may require blockade of multiple inhibitory receptors (PD-1, CTLA-4, and LAG-3) [38]. Thus, our work indicates that p38 blockade could reverse (or prevent) T cell exhaustion in the TME, leading to enhanced T cell mediated anti-tumor responses.

With respect to other states of CD8^+^ T cell differentiation, blockade of p38α increased the frequency of CD62L^high^CD44^+^ CD8^+^ T cell population in both TdLNs and spleen. This subset of CD8^+^ T cells exhibits a stem-like memory phenotype, also known as central-memory T cells (T_CM_), and elicits superior anti-tumor immunity in the context of adoptive cell transfer [59, 60]. These cells express lower levels of effector molecules (*Prf1, Gzmb*) compared to the effector-memory subset but engraft more efficiently and persist longer in patients [61]. Our *in vitro* data indicate that p38 may directly control expression of CD44 and CD62L on CD8^+^ T cells (**Fig. 7c**). However, p38 blockade showed only marginal inhibitory effects on T cell proliferation in *in vitro* assays. The increased levels of CD62L on CD8^+^ T cells align with the role of CD62L in T cell homing to areas of lymphoid tissues [61]. In agreement with our findings, a recent study identified p38 as an important regulator of CD62L^+^ CD8^+^ T cells in mouse models [62]. Thus, pharmacological inhibition of p38 may exhibit a direct effect on CD8^+^ T cell phenotype, in addition to the indirect effects elicited by tumor-specific p38 signaling discussed above.

Major effects of p38 blockade were observed on suppressive myeloid cells. Blockade of p38 by both pharmacological and genetic approaches suppressed tumor-induced expansion of TAMs and MDSCs (**Fig. 5**). Transcriptomic analysis of the TME showed that p38 blockade reduced the expression of markers for granulocytic (PMN) and monocytic populations and reduced the expression of chemokine receptors by myeloid cells. Furthermore, p38 blockade decreased the levels of monocytes and granulocytes in circulation and reduced the expression of multiple immune suppressive markers (*e.g.*, *Vegfa, Tgf*β*1, Arg1*) in splenic granulocytic subsets. These results imply that p38 controls the mechanisms governing tumor-induced expansion and recruitment of immune suppressive myeloid cells, which, in turn, induce exhaustion of T cells. In support of this indirect effect of p38, prior work showed that p38i did not block the generation of immune suppressive MDSCs *in vitro* [11].

Investigations into how p38 may regulate the expansion and recruitment of TAMs and MDSCs showed that p38α controls the tumor cell expression of cytokines and chemokines (e.g., CCL2, CXCL1, IL6, GM-CSF, G-CSF). These signals can stimulate the expansion and recruitment of myeloid populations, such as PMN-MDSCs and TAMs [35, 54, 57, 63]. Previous studies have shown that tumor-derived CCL2 promotes the recruitment of TAMs and MDSCs [64, 65], while CXCL1 promotes the recruitment of MDSCs [66].

Importantly, our work revealed an unrecognized role of p38α in regulating tumor-derived exosomes with a chemotactic capacity. Exosomes are cell-secreted vesicles (30-150nm diameter) that mediate cell to cell communication by carrying signaling molecules, which are taken up by other cells and influence the phenotype of those cells [67]. We found that p38 blockade (by p38i or p38KO) reduced the amounts and chemotactic activity of exosomes secreted by human and mouse tumor cells, indicating that p38α regulates the production and the content of exosomes. A prior study showed that exosomes secreted by mouse TNBC cell lines (such as 4T1) are mainly distributed to the lungs and promote metastatic colonization in the lungs [68]. 4T1-derived exosomes are primarily taken up by myeloid cells in the lungs (40-60%) and spleen (20-30%) and markedly increased the frequency of PMN-MDSCs [68]. Mechanistically, tumor-derived exosomes may carry chemokines (i.e., CCL2) that, by interacting with cognate receptors (i.e., CCR2), promote the uptake of exosomes by myeloid cells at metastatic sites [69]. Further, 4T1-derived exosomes can also carry matrix proteins, such as fibronectin [70] that exhibit chemo-attraction for myeloid cells [71]. In support of this notion, p38α controls the expression of CCL2 and fibronectin in mouse and human TNBC cell lines, including 4T1 and MDA-MB-231 [11, 72]. Taken together, these observations support a critical role for p38α in the production of tumor-derived exosomes and signaling molecules (cytokines/chemokines) that can mediate the expansion and recruitment of TAMs and MDSCs.

Altogether, our study demonstrates that pharmacological targeting of p38α may improve the antitumor immune response by restraining multiple integral elements of the immune suppressive TME. These findings implicate p38α as a druggable therapeutic target and that pharmacological blockade of p38α can be combined with immune-based therapies, such as immune checkpoint inhibitors, to improve therapeutic responses and patient outcomes.

## LIST OF ABBREVIATIONS

MDSCs: myeloid-derived suppressive cells;
TAMs: tumor-associated macrophages;
TME: tumor microenvironment;
TNBC: triple-negative breast cancer;
MBC: metastatic breast cancer.

## ACKNOWLEDGEMENTS

We gratefully acknowledge the generous help from Flow and Image Cytometry Facility, the Genomics Shared Resource, Gene Targeting and Transgenic Resource (GTTR) and Comparative Oncology Shared Resource. We thank Mohammed Alqarni and Anand Sharda for help in the study, the Deanship of Scientific Research at Northern Border University, Arar; We thank Dr. Aimee Stablewski for help with CRISPR/Cas9, Dr. Kevin Eng, Dr. Michalis Mastri and Dr. Mark Long for help with bioinformatic analysis.

## AUTHOR CONTRIBUTIONS

A.V.B. and S.I.A. developed the study; A.V.B., S.I.A., J.B., and S.O. designed the experiments; P.R., R.Z., Y.G., M.A., M.L., J.Z.,C.J., A.V.B, and B.M. performed the experiments and analyzed the data; P.R., A.V.B., J.B., and S.O., analyzed and interpreted the data; A.V.B., P.R., S.I.A., J.B., and S.O. contributed to preparing and writing the paper.

## DATA AVAILABILITY STATEMENT

The data generated in this study are available upon request from the corresponding author. The scRNAseq raw data have been deposited at NCBI GEO site access number GSE282734.

## ETHIC STATEMENT

A study protocol and guidelines approved by the Institute Animal Care and Use Committee (IACUC). The facility is certified by the American Association for Accreditation of Laboratory Animal Care (AAALAC) and in accordance with current regulation and standards of the US Department of Agriculture and the US Department of Health and Human Services.

