## Supplementary figures for "p38 blockade reverses the immune suppressive tumor microenvironment in metastatic breast cancer"

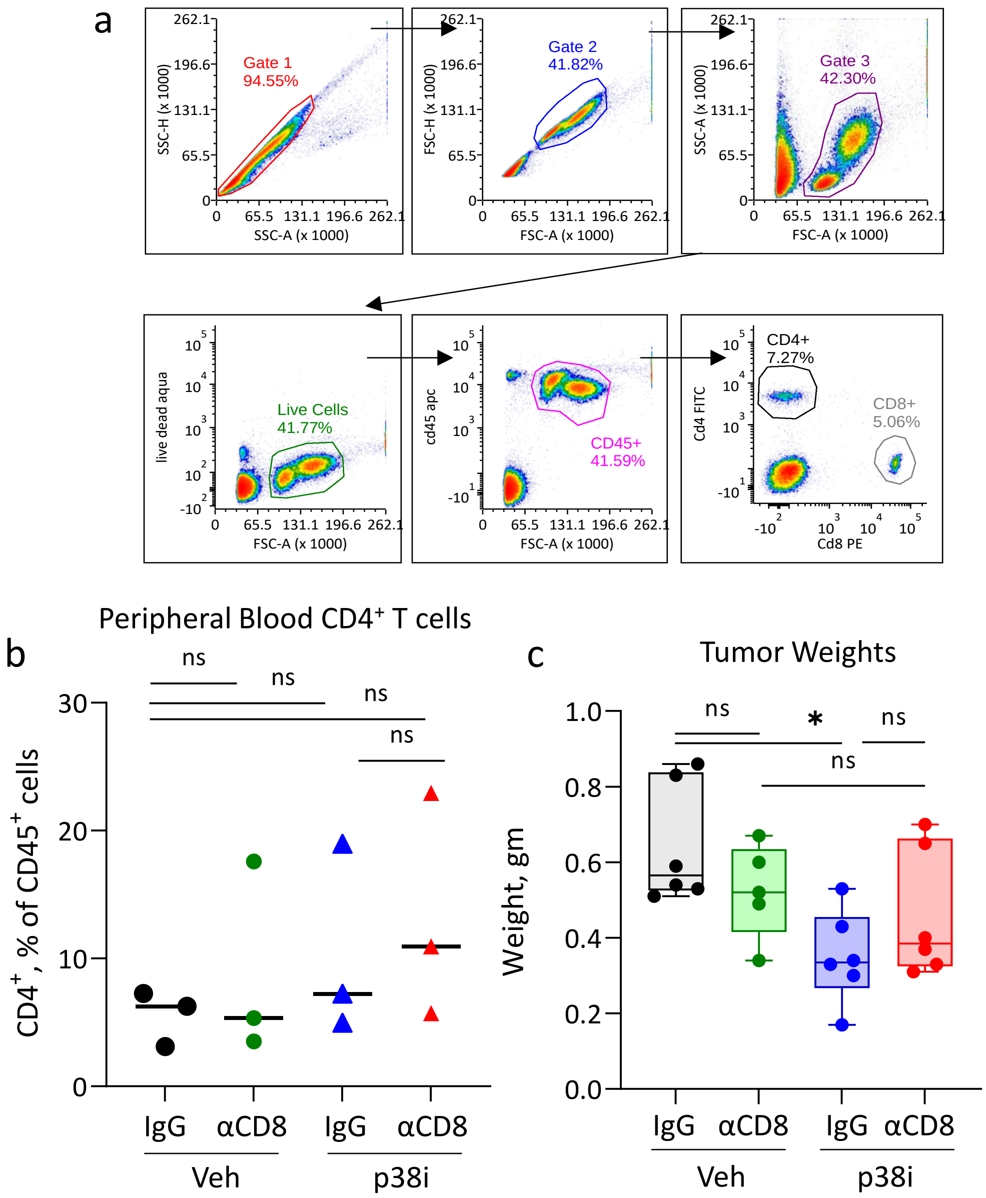


**Supplementary Fig. 1.** p38 blockade reduced 4T1 lung metastasis in a CD8⁺ T cell dependent manner. (**a**) Gating strategy for flow cytometric evaluation of CD8⁺ T cells in mouse peripheral blood. (**b**) Flow cytometric evaluation of CD4⁺ T cell levels in peripheral blood of mice (n=3/group). (**c**) Tumor weights at endpoint. Statistical analysis was done by one-way ANOVA (*, p<0.05; ns, no significance).

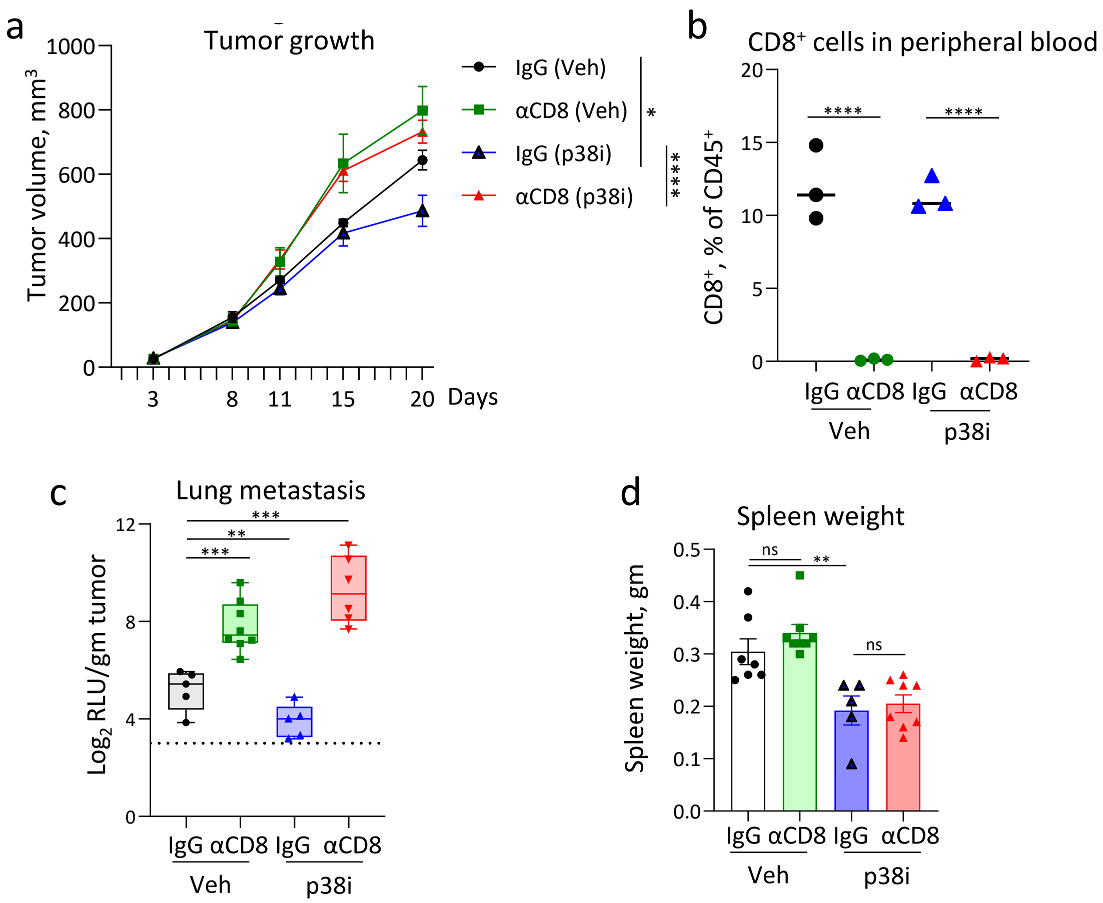


**Supplementary Fig. 2. CD8$^+$ T cells contribute to anti-metastatic activity of p38i**. Mouse mammary carcinoma 4T1 cells were implanted into mammary fat pads of BALB/c mice. On day 3, mice received i.p. injections of antibody to CD8 or IgG control. p38i was given daily starting on day 4. Tumor volumes were measured 2 times per week. (**a**) Tumor volume change over time, comparisons were done by one-way ANOVA. (**b**) Levels of CD8$^+$ T cells in peripheral blood were measured by flow cytometry on day 6 post tumor implantation. Comparisons were done by one-way ANOVA. (**c**) Luciferase activity in whole cell extracts from the lungs in tumor-bearing mice, n=8/group. Values are shown in log2 values normalized to average tumor weight. RLU - Relative Luminescence Units. Comparisons were done by one-way ANOVA. (**d**) Spleen weights at endpoint. Comparisons were done by one-way ANOVA. *, $p<0.05$; **, $p<0.01$; ***, $p<0.001$; ****, $p<0.0001$; ns, non-significant.

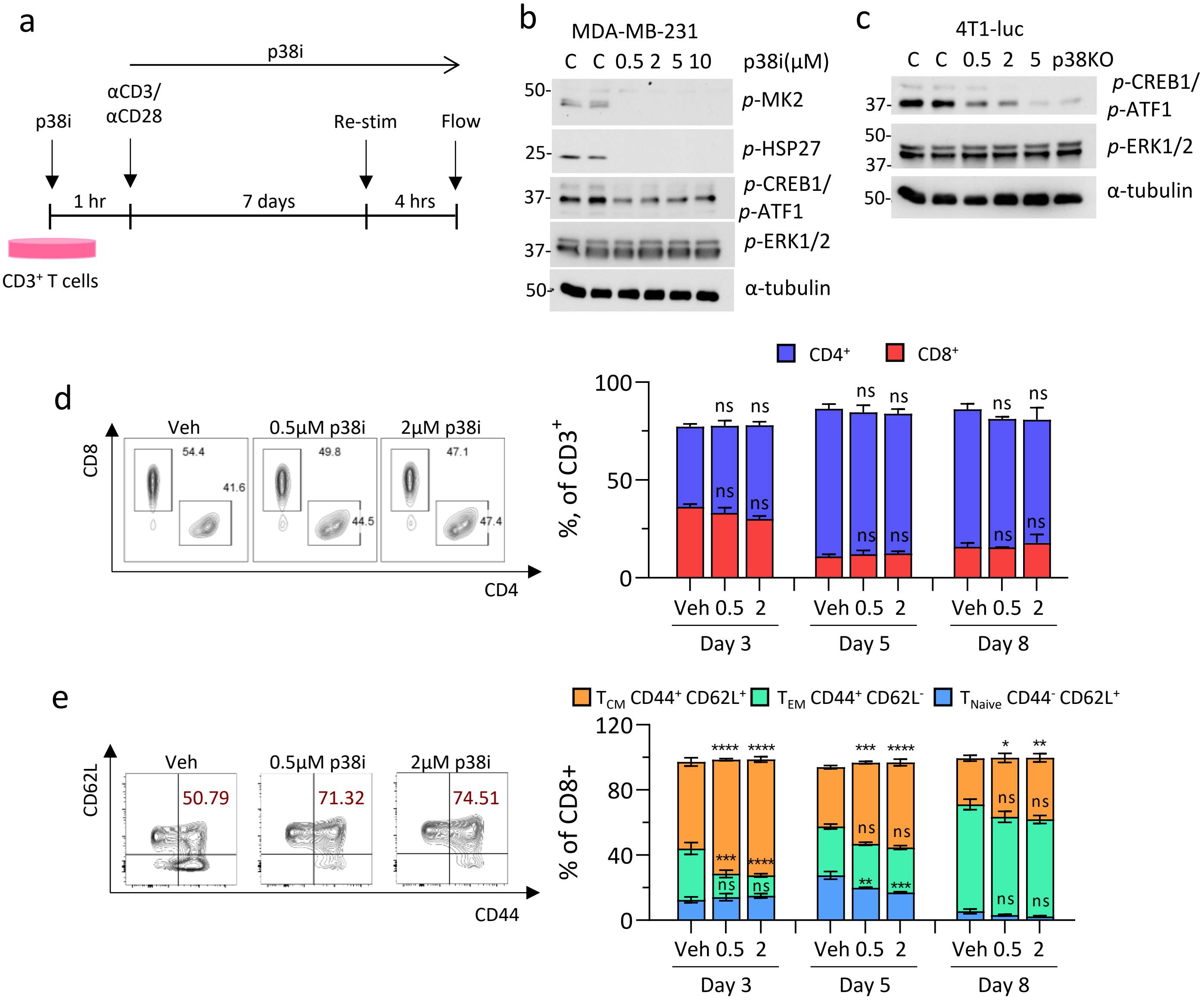


**Supplementary Fig. 3. p38 blockade has little effects on T cells cultured *ex vivo.*** (**a**) Scheme of in vitro T cell activation assay. (**b**) & (**c**) MDA-MB-231 and 4T1-luc cells were treated with varying doses of p38i for 6hrs. Immunoblotting was performed on the lysates and probed with antibodies against p-MK2 (T-222), p-HSP27, p-CREB1/p-ATF1 (S133), p-ERK1/p-ERK2 (T202/Y204) and alpha-tubulin. 4T1-luc p38KO1 lysates were run under the same conditions. (**d**) Percentage of CD4+ and CD8+ T cells at each time point. (**e**) Percentage of CD8+ that are CD44+ CD62L+ ($T_{CM}$ like), CD44- CD62L+ ($T_{naive}$), and CD44+ CD62L- ($T_{EM}$ like). Flow plots are representative of Day 3. (Continued on next page)

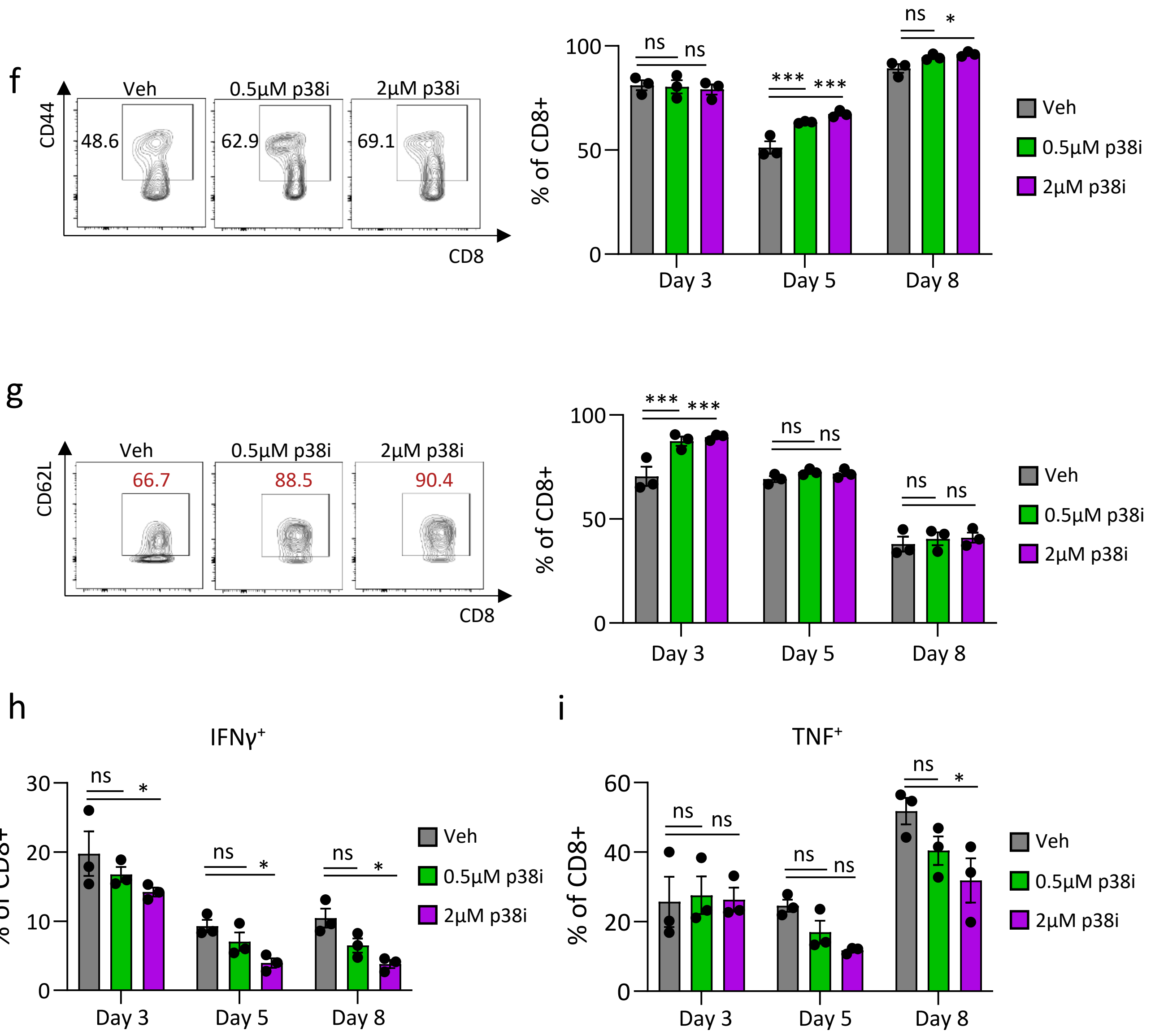


**Supplementary Fig. 3 (Continued).** (**f**) Percentage of CD8+ T cells expressing CD44. Flow plots are representative of Day 3. (**g**) Percentage of CD8+ T cells expressing CD62L. Flow plots are representative of Day 3. (**h**) Percentage of CD8+ T cells expressing IFNγ. (**i**) Percentage of CD8+ T cells expressing TNF. For (**d-i**), each of the treated groups was compared to its own vehicle group by two-way ANOVA with Tukey's multiple comparisons test. * $p < 0.05$, ** $p<0.01$, *** $p<0.001$, **** $p<0.0001$ and ns = non-significant. (Gating strategy on next page).

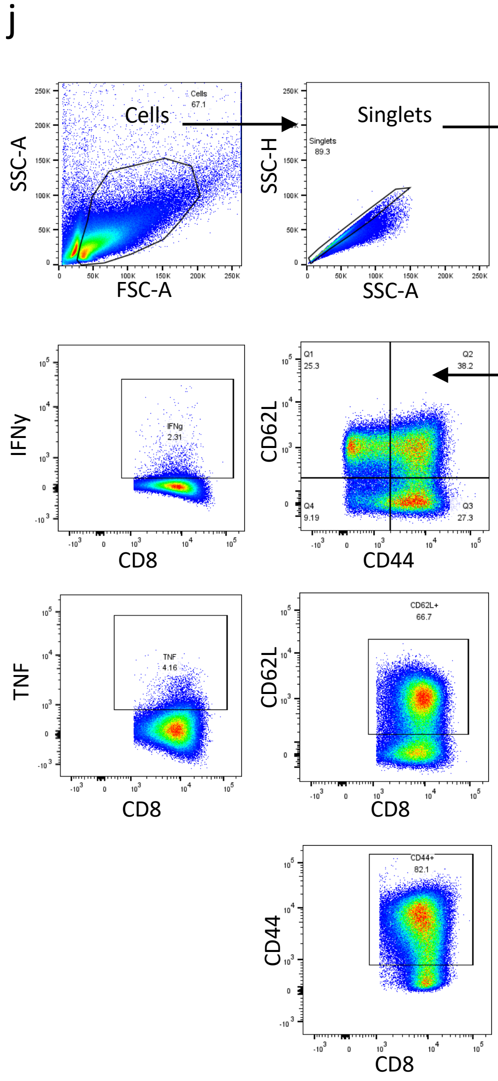


**Supplementary Fig. 3 (Continued).** (j) Gating strategy for *in vitro* T cell activation assay.

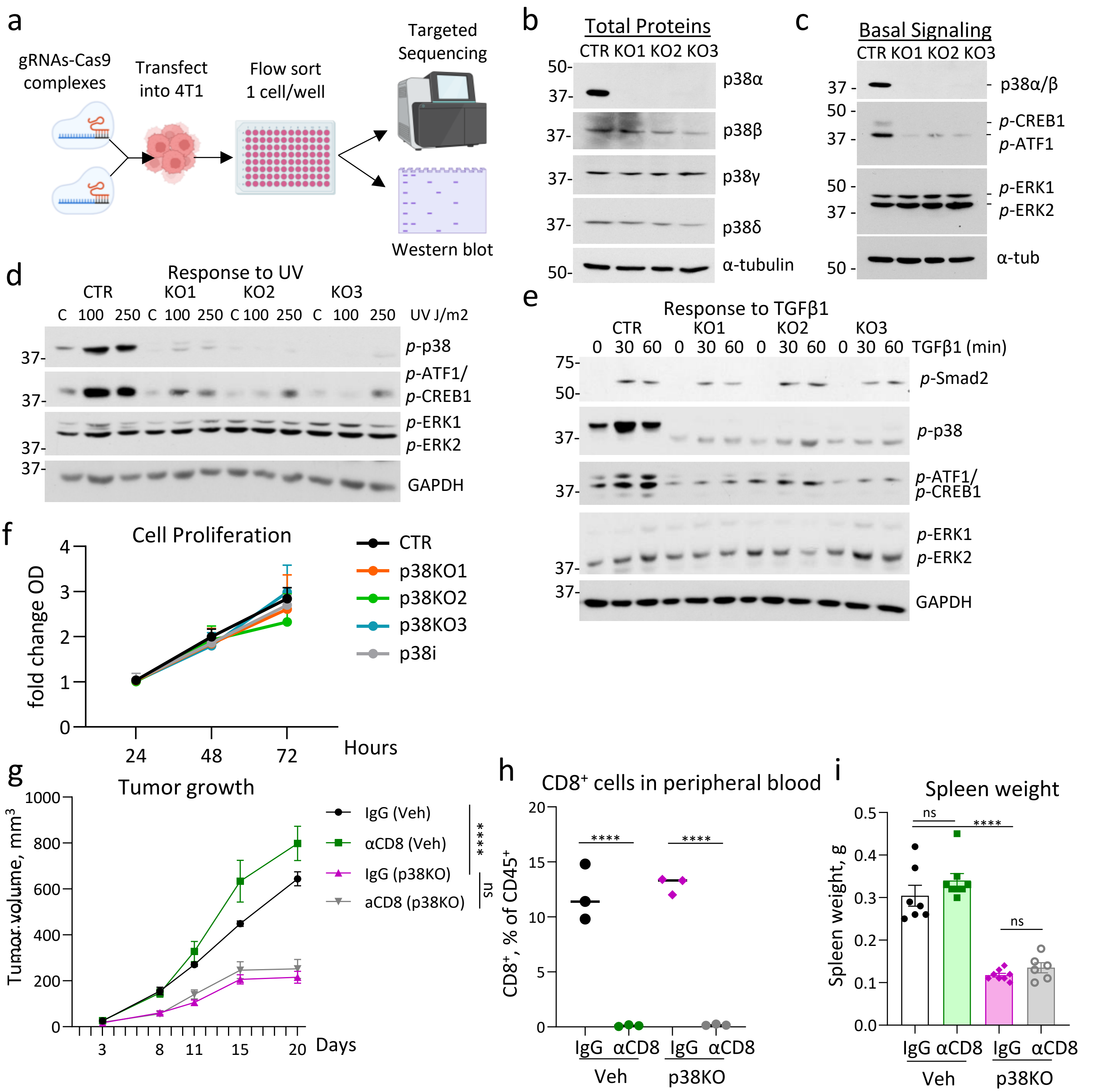


**Supplementary Fig. 4.** Generation and characterization of 4T1 p38KO clones. (**a**) Workflow. (**b**) Immunoblot of lysates of 4T1-luc control (CTR) and p38KO cells probed for p38α, p38β, p38γ, p38δ and α-tubulin. (**c**) Immunoblot of 4T1-luc (CTR) and p38KO cells probed for p38α/β, p-CREB1/p-ATF1 (S133), p-ERK1/p-ERK2 (T202/Y204), and tubulin as loading control. (**d**) *In vitro* growth curves of 4T1 (control) and p38KO cells. Cells were treated with vehicle control (DMSO) or p38i for 24, 48 or 72h. Graph shows fold change compared to CTR and represents one of 3 biological replicates. (**e**) Cells were treated with TGFβ1 for 0, 30 or 60 mins. Immunoblots were probed for p-SMAD2 (S432/T434), p-p38, p-CREB1/ATF1, p-ERK1/2 (T202/Y204), and GAPDH. (**f**) Cells were subjected to 0, 100 or 250J/m$^2$ UV radiation. Lysates were probed for p-p38, p-CREB1/ATF1, p-ERK1/ERK2 and GAPDH. The immunoblots are representative of 3 biological repeats. (**g**) 4T1 or 4T1-p38KO1 cells were implanted into mammary fat pads of BALB/c mice. On days 3, 10, and 17 mice received i.p. injections of antibody to CD8 or IgG control. Tumor volumes were measured 2 times per week. (**h**) Levels of CD8$^+$ T cells in peripheral blood were measured by flow cytometry on day 6 post tumor implantation. (**i**) Spleen weights at the endpoint. Comparisons were done by one-way ANOVA. *, p<0.05; **, p<0.01; ***, p<0.001; ****, p<0.0001; ns, non-significant.

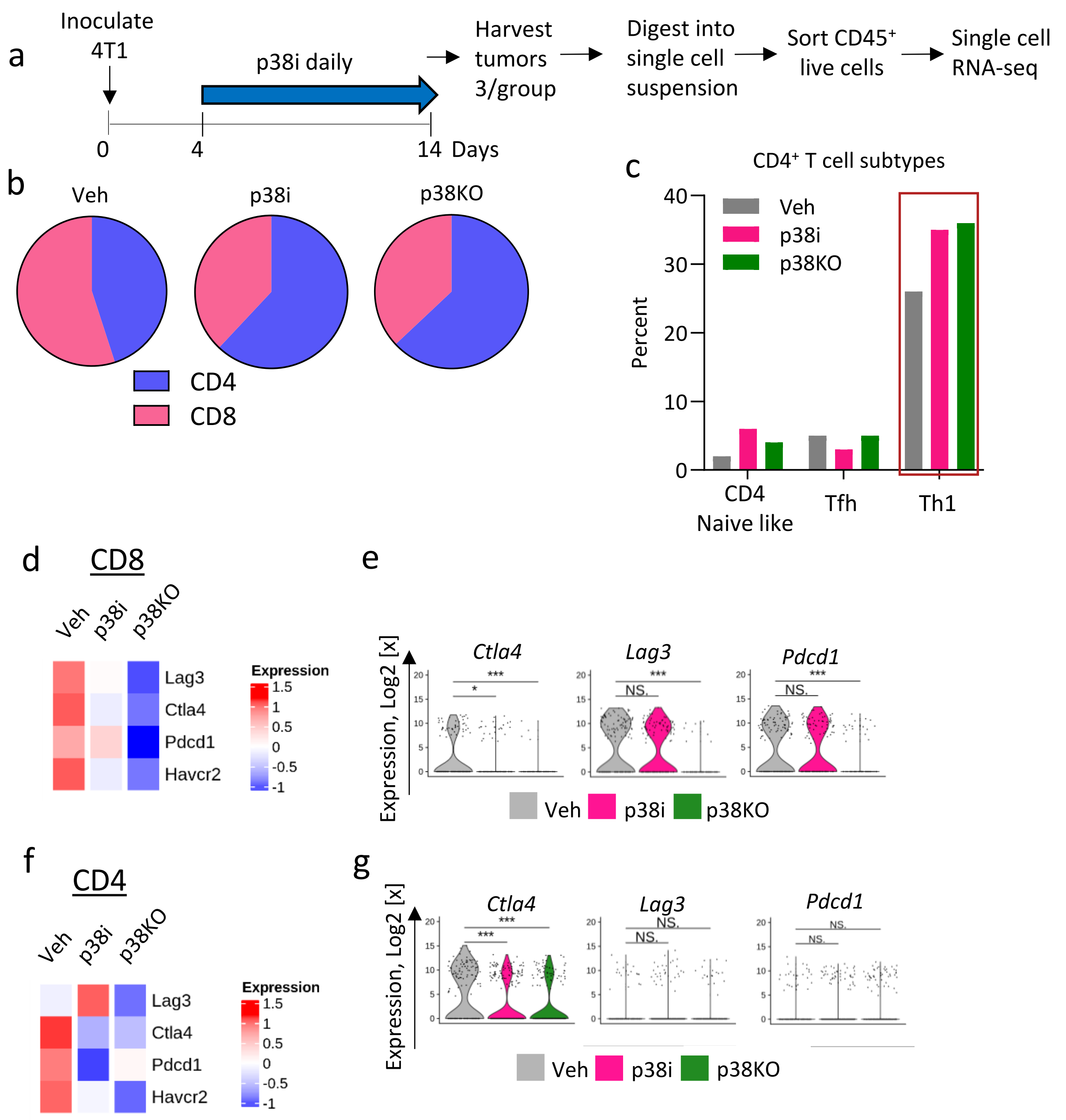


**Supplementary Fig. 5. Inactivation of p38 alters the tumor immune transcriptomic landscape**. (**a**) Scheme of the single cell RNA-seq study. (**b**) Proportion of $CD4^+$ and $CD8^+$ T cells identified by ProjecTILs R package in each group. (**c**) Percentage of $CD4^+$ T cell sub-populations identified in each group by ProjecTILs R package. (**d**) Heatmap showing scaled average expression levels of genes in CD8+ T cells from vehicle control or p38i treated or p38KO tumors, measured in log2 counts per million. (**e**) Violin plots showing expression of specific genes measured in log2 counts per million, in the CD8+ T cells from vehicle control or p38i treated or p38KO tumors. (**f**) Heatmap showing scaled average expression levels of genes in CD4+ T cells from vehicle control or p38i treated or p38KO tumors, measured in log2 counts per million. (**g**) Violin plots showing expression of specific genes measured in log2 counts per million, in the CD4+ T cells from vehicle control or p38i treated or p38KO tumors. Comparisons are done by t-test.

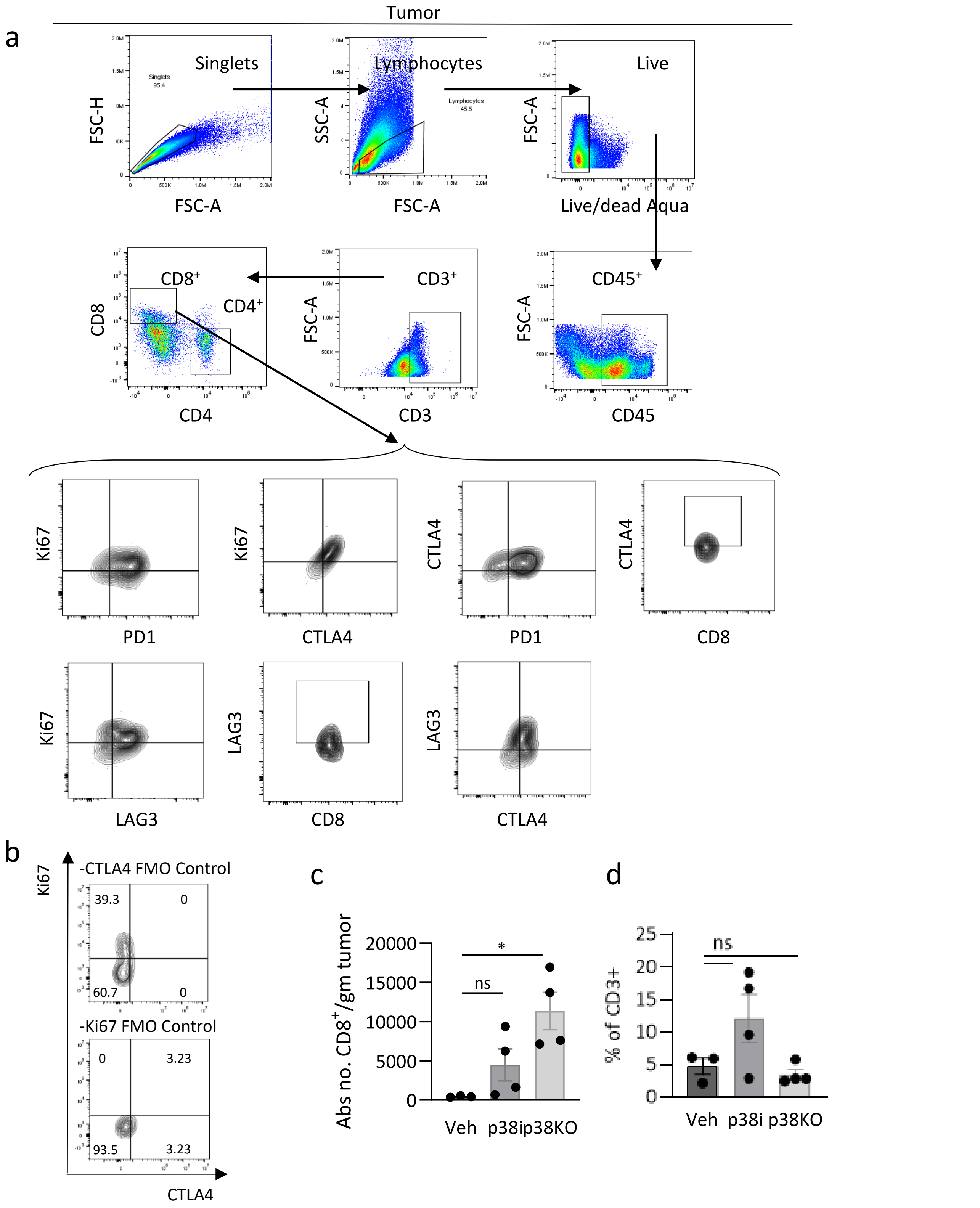


**Supplementary Fig. 6. Immunophenotyping of T cells in the tumor treated with p38i or p38 deleted.** (**a**) Gating strategy for T cells in the tumor. (**b**) FMO controls for CD8+ T cells in the tumor expressing CTLA4 and Ki67. Data are representative of 3 individual experiments and are plotted as mean +- SEM for n=3-4 mice/group. Statistical analysis was performed by two-tailed unpaired Student's t-test. * p < 0.05, ** p<0.01, *** p<0.001, **** p<0.0001 and ns = non-significant. (**c**) Number of CD8+ T cells per gram tumor. (**d**) Percentage of CD8+ T cells in CD3 population. Comparisons were made using two-way ANOVA. *, p<0.05; **, p<0.01; ***, p<0.001; ****, p<0.0001; ns, non-significant.

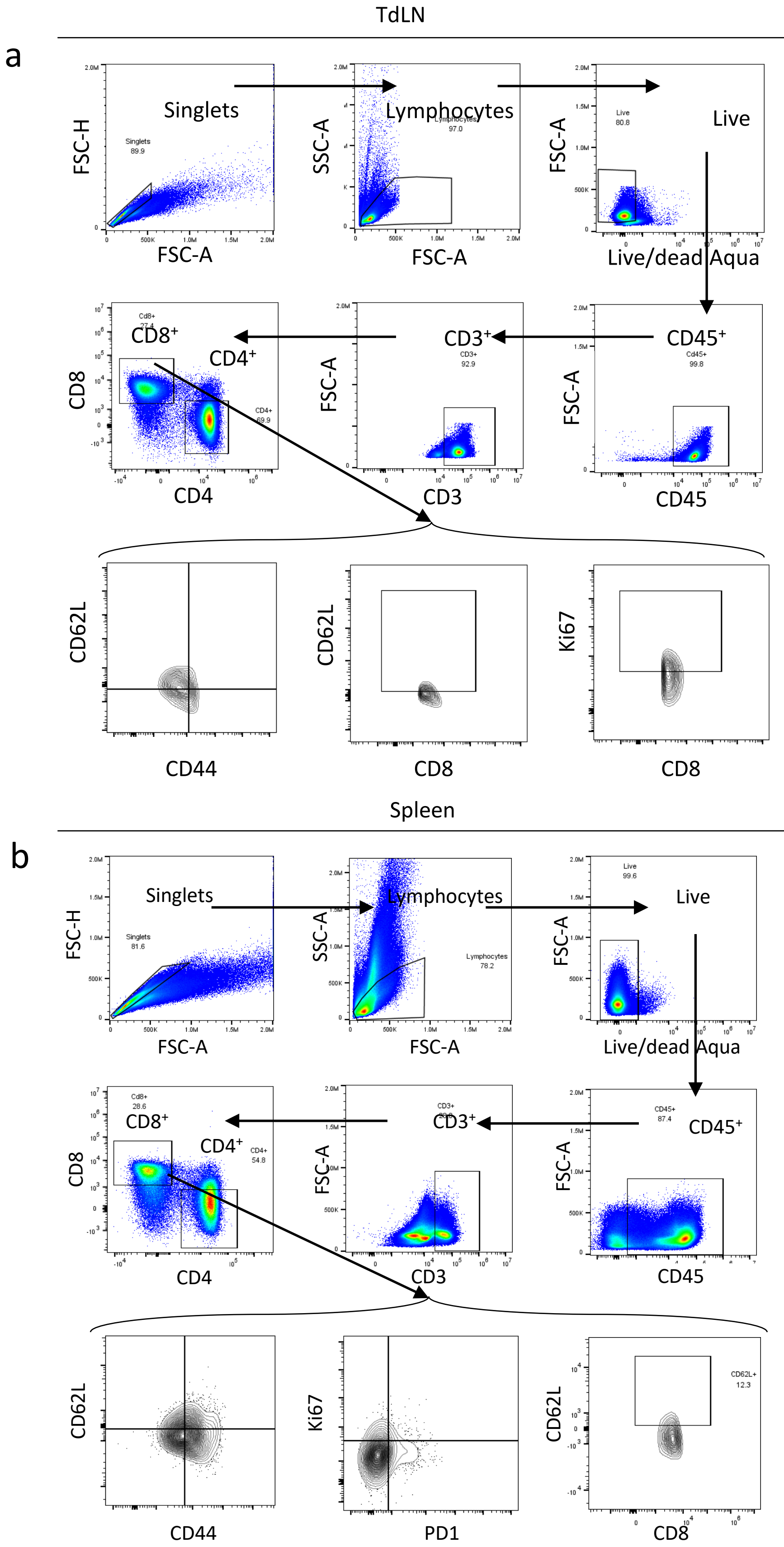


**Supplementary Fig. 7. p38 blockade reduces T cell exhaustion: the analysis of TdLNs and spleens.** (**a**) Gating strategy for T cells in the tumor draining lymph node. (**b**) Gating strategy for T cells in the spleen. (Continued on next page)

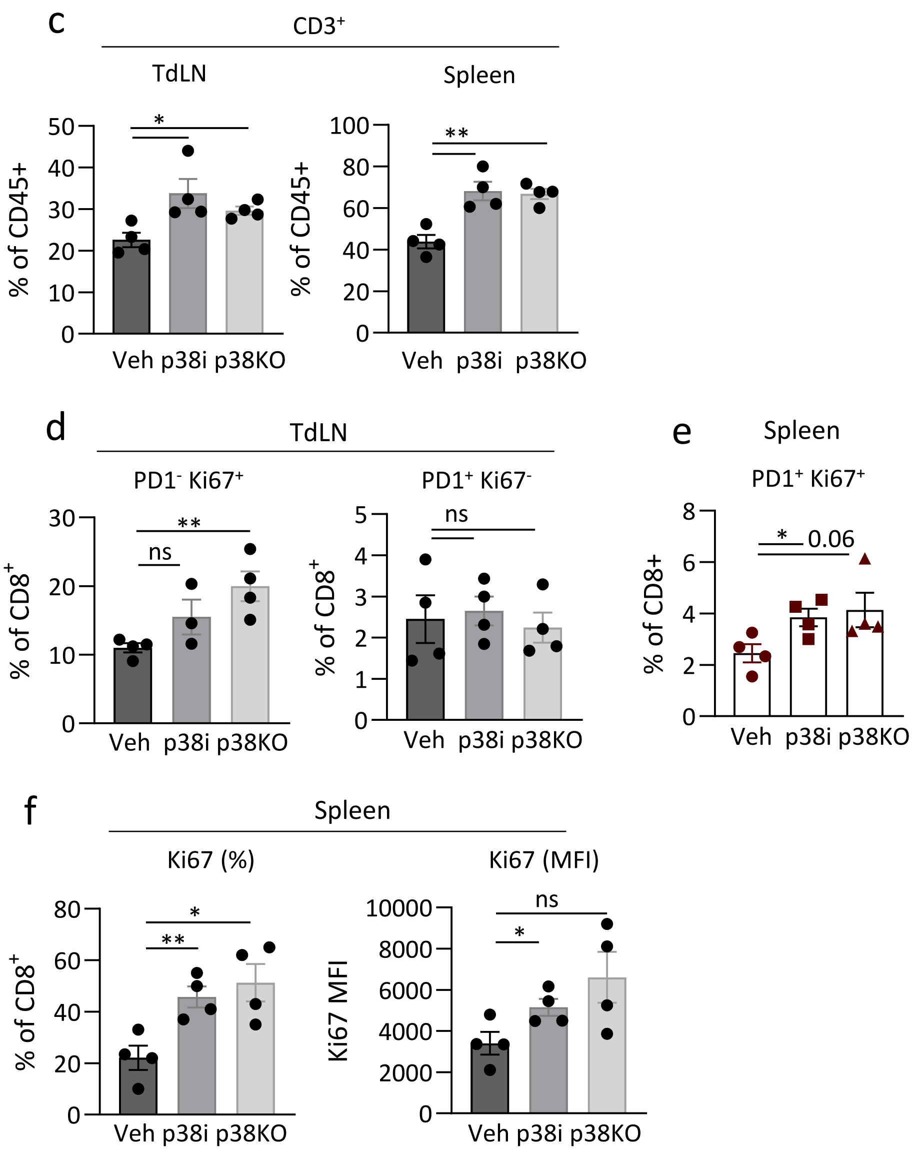


**Supplementary Fig. 7 (continued).** (**c**) Percentage of $CD3^+$ T cells in the TdLN and spleen. (**d**) Percentage of $CD8^+$ T cells in the TDLNs that are $PD1^+ Ki67^-$ and $PD1^- Ki67^+$. (**e**) Percentage of $CD8^+$ T cells in spleen expressing $PD1^+ Ki67^+$. (**f**) Percentage of $CD8^+$ T cells in spleen expressing $Ki67^+$ by MFI and percentage. Data are representative of 3 individual experiments and are plotted as mean +- SEM for n=3-4 mice/group. Statistical analysis was performed by two-tailed unpaired Student's t-test. * $p < 0.05$, ** $p<0.01$, *** $p<0.001$, **** $p<0.0001$ and ns = non-significant. (Continued on next page).

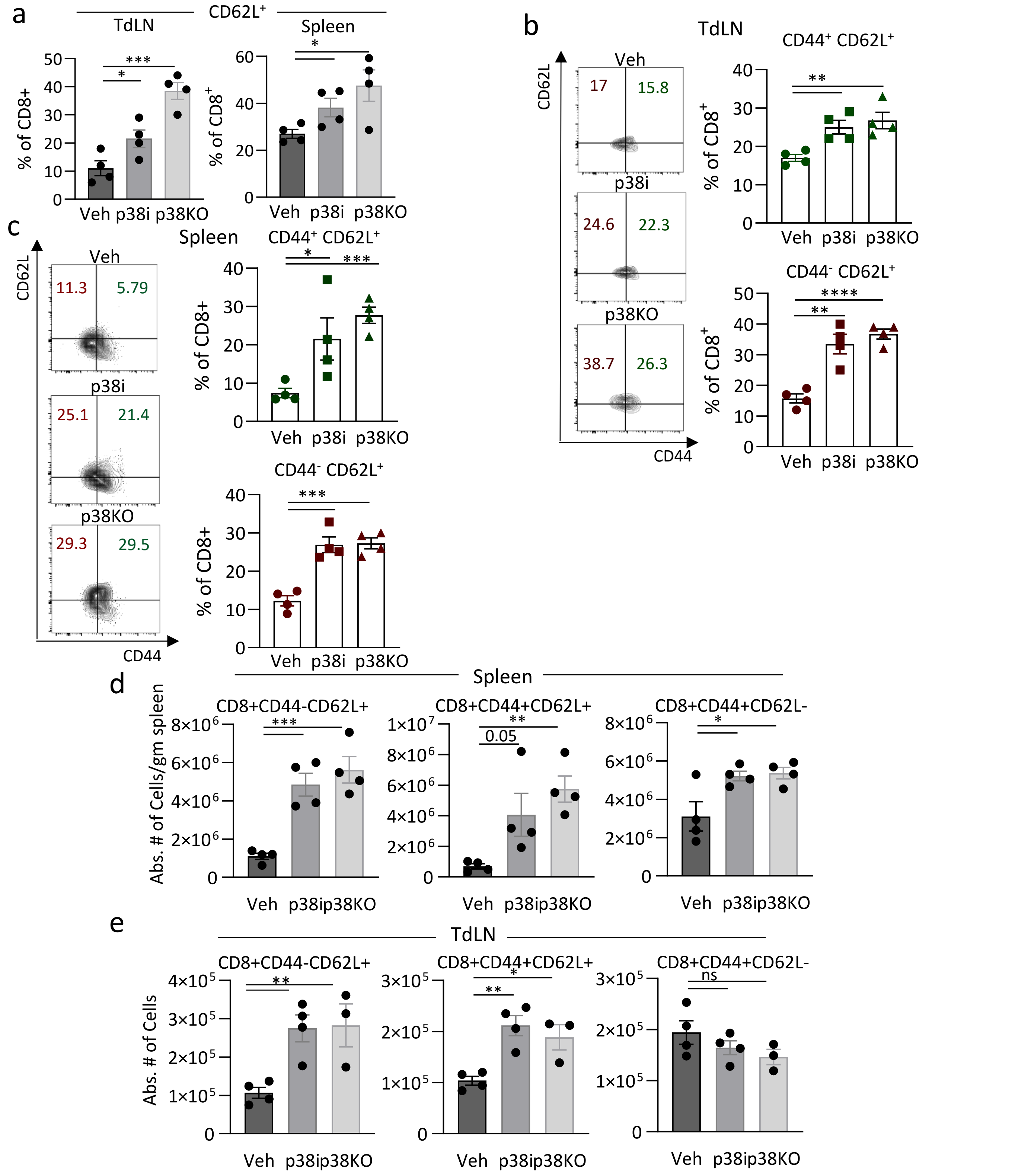


**Supplementary Fig. 8. p38 blockade enhanced CD62L⁺ population.** (**a**) Percentage of CD62L⁺ T cells in the TdLN and spleen. (**b**) Percentage of cells that are CD44⁻ CD62L⁺ and CD44⁺ CD62L⁺ within the CD8⁺ T cells in the TdLNs. (**c**) Percentage of cells that are CD44⁻ CD62L⁺ and CD44⁺ CD62L⁺ within the CD8⁺ T cells in the spleen. Data are representative of 3 individual experiments and are plotted as mean +- SEM for n=3-4 mice/group. (**d**) Absolute # of CD8+ cells in the spleen: $T_{naive}$ (CD44- CD62L+), $T_{CM}$ (CD44+ CD62L+) and $T_{EM}$ (CD44+ CD62L-). (**e**) Absolute # of CD8+ cells in the TdLNs: $T_{naive}$ (CD44- CD62L+), $T_{CM}$ (CD44+ CD62L+) and $T_{EM}$ (CD44+ CD62L-). Comparisons by Student's t-test. * $p < 0.05$, ** $p<0.01$, *** $p<0.001$, **** $p<0.0001$ and ns = non-significant.

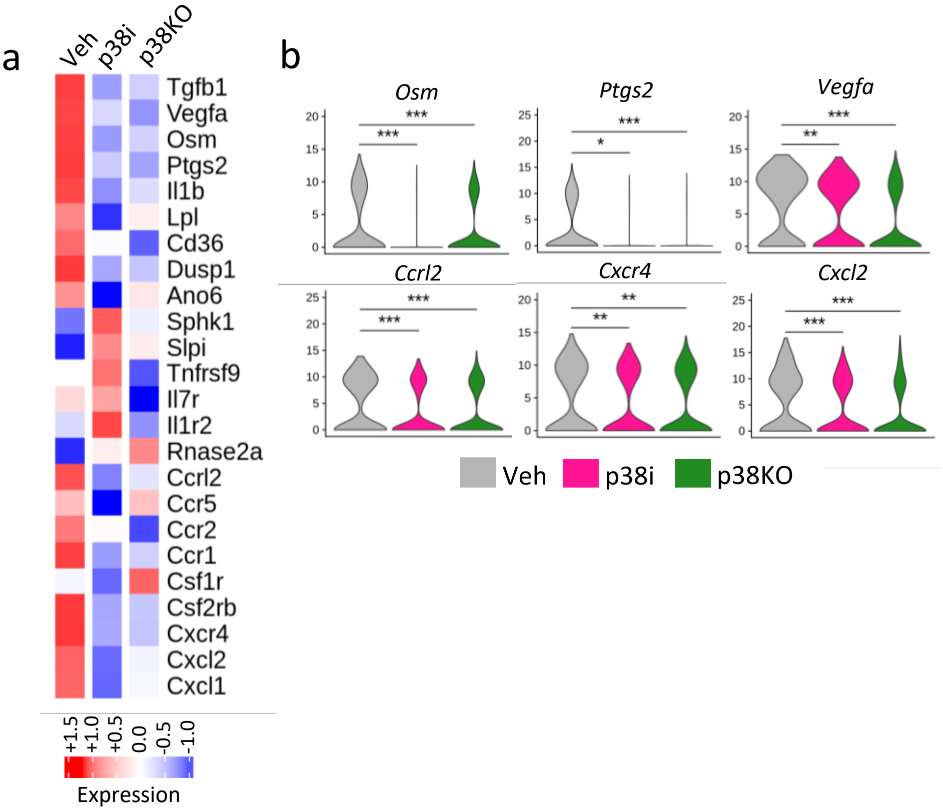


**Supplementary Fig. 9. Changes to tumor macrophage transcriptome caused by p38 inactivation.** (**a**) Heatmap of scRNAseq data showing scaled average expression levels of genes in the macrophages from vehicle control or p38i treated or p38KO tumors measured in log2 counts per million. (**b**) Violin plots showing distribution of expression levels of specific genes, measured in log2 counts per million, in the macrophages from vehicle control or p38i treated or p38KO tumors. Statistical analysis was done by t-test, * p < 0.05, ** p<0.01, *** p<0.001, **** p<0.0001, ns = non-significant.

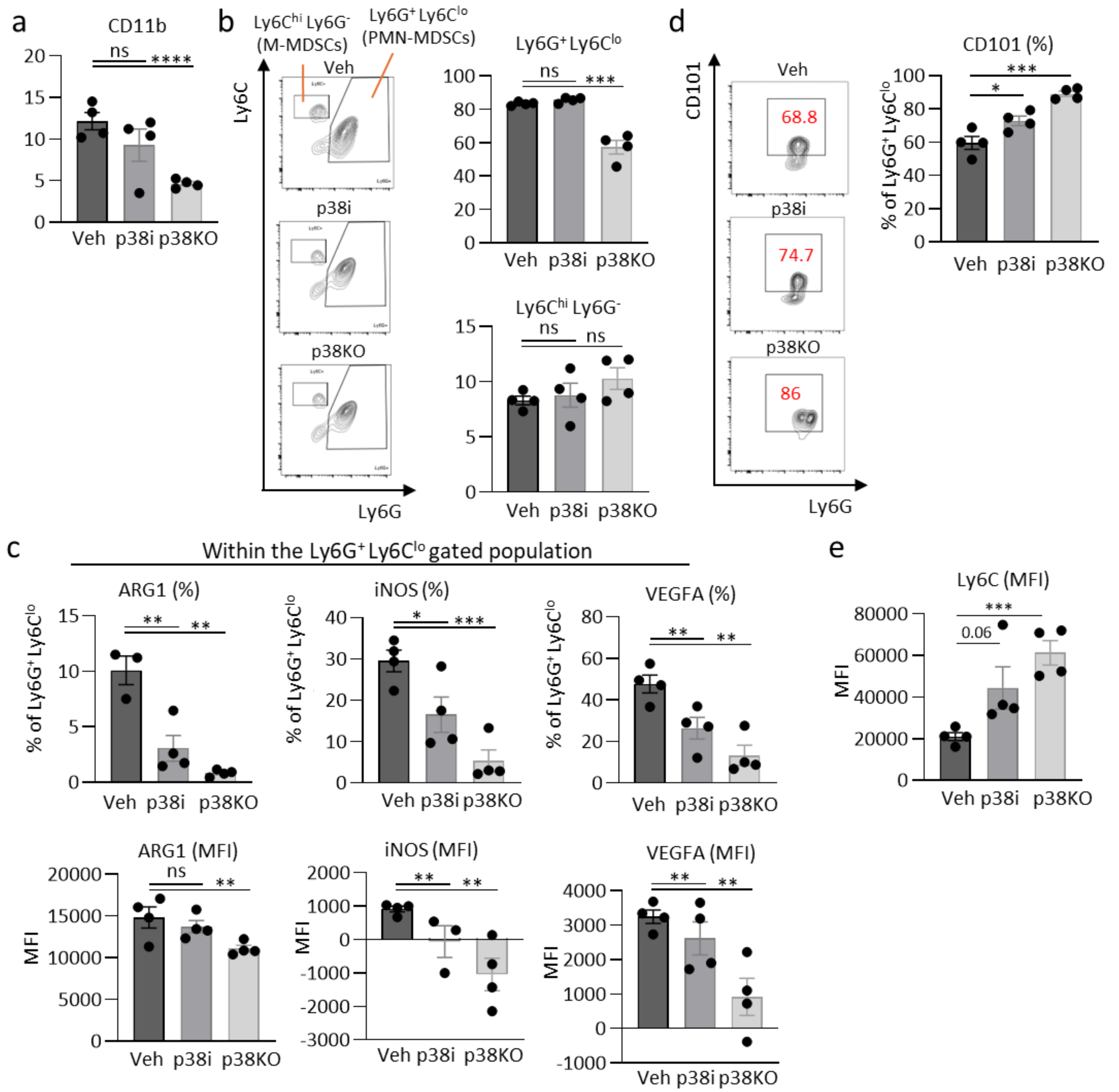


**Supplementary Fig. 10. Blockade of p38 alters the phenotype of MDSCs in the spleen.** 4T1 p38 wild type (WT, control) and p38KO cells were implanted into mammary fat pads of BALB/c mice. Mice bearing 4T1 p38WT tumors were treated with vehicle (PBS) or p38i (Ralimetinib). At the experimental endpoint (day 14 post tumor inoculation), spleens were harvested and processed for flow cytometry. (**a**) Percentage of $CD11b^{+}$ cells in the spleen. (**b**) Percentage of $Ly6G^{+}$ $Ly6C^{lo}$ or $Ly6C^{hi}$ $Ly6G^{-}$ cells in the spleen. (**c**) Expression of ARG1, iNOS or VEGFA (percentage and MFI) by $Ly6G^{+}$ $Ly6C^{lo}$ cells in the spleen. (**d**) Expression of CD101 (percentage and MFI) by $Ly6G^{+}$ $Ly6C^{lo}$ cells in the spleen. (**e**) Expression of Ly6C (MFI) by $Ly6G^{+}$ $Ly6C^{lo}$ cells in the spleen. Data are presented as mean +- SEM for 2 independent experiments. Statistical analysis was done by t-test (*, p<0.05; **, p<0.01; ***, p<0.001; ****, p<0.0001).

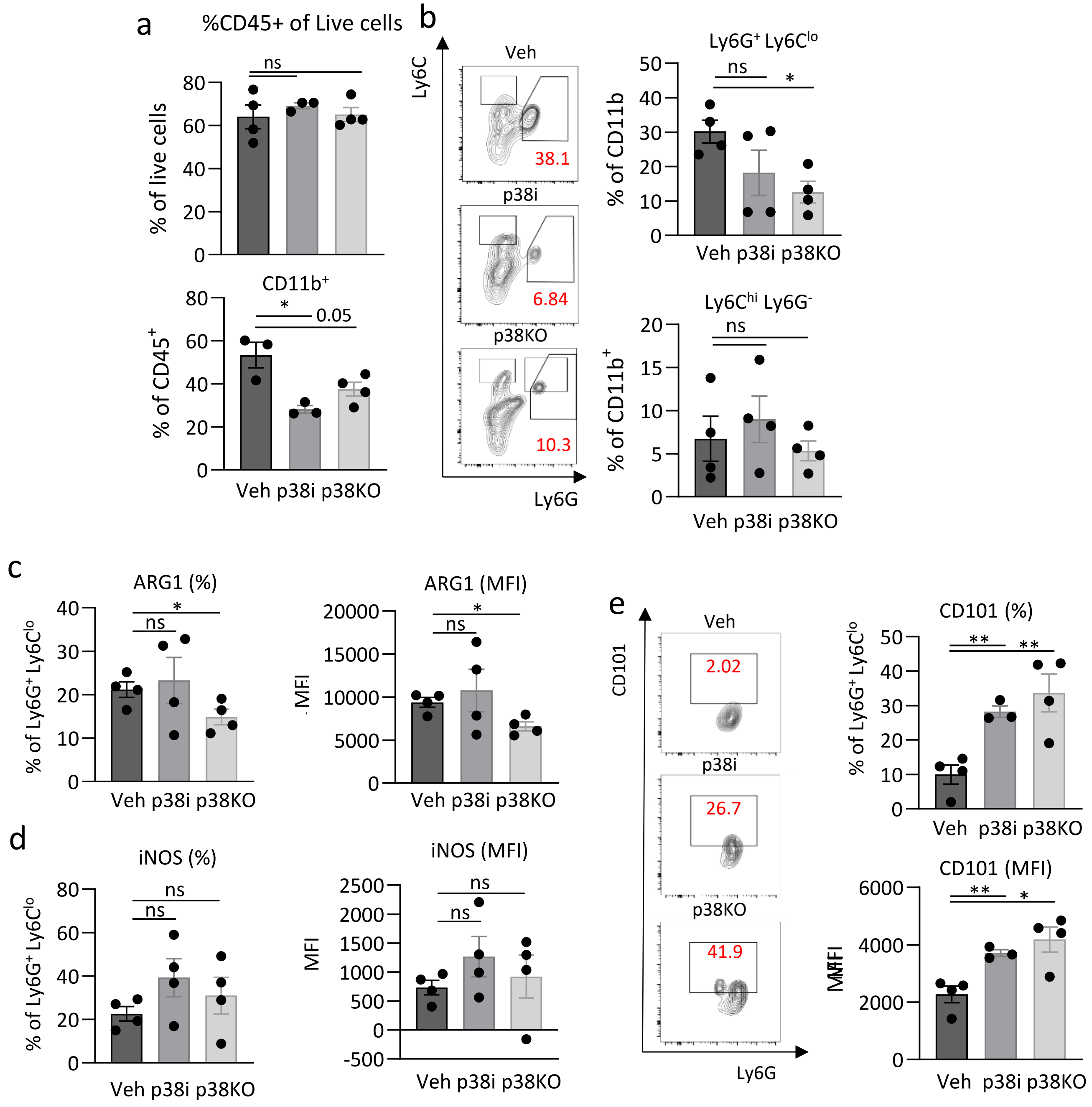


**Supplementary Fig. 11. Inactivation of p38 reduces myeloid cell accumulation in the tumor.** (**a**) Percentage of CD11b$^+$ cells in the tumor. (**b**) Percentage of Ly6G$^+$ Ly6C$^{lo}$ and Ly6C$^{hi}$ Ly6G$^-$ cells in the tumor. (**c**) Expression of ARG1 (percentage and MFI) by Ly6G$^+$ Ly6C$^{lo}$ cells in the tumor. (**d**) Expression of iNOS (percentage and MFI) by Ly6G$^+$ Ly6C$^{lo}$ cells in the tumor. (**e**) Expression of CD101 (percentage and MFI) by Ly6G$^+$ Ly6C$^{lo}$ cells in the tumor. Data are presented as mean +- SEM for 2 independent experiments. Comparisons are done by t-test (*, p<0.05; **, p<0.01; ***, p<0.001; ****, p<0.0001).

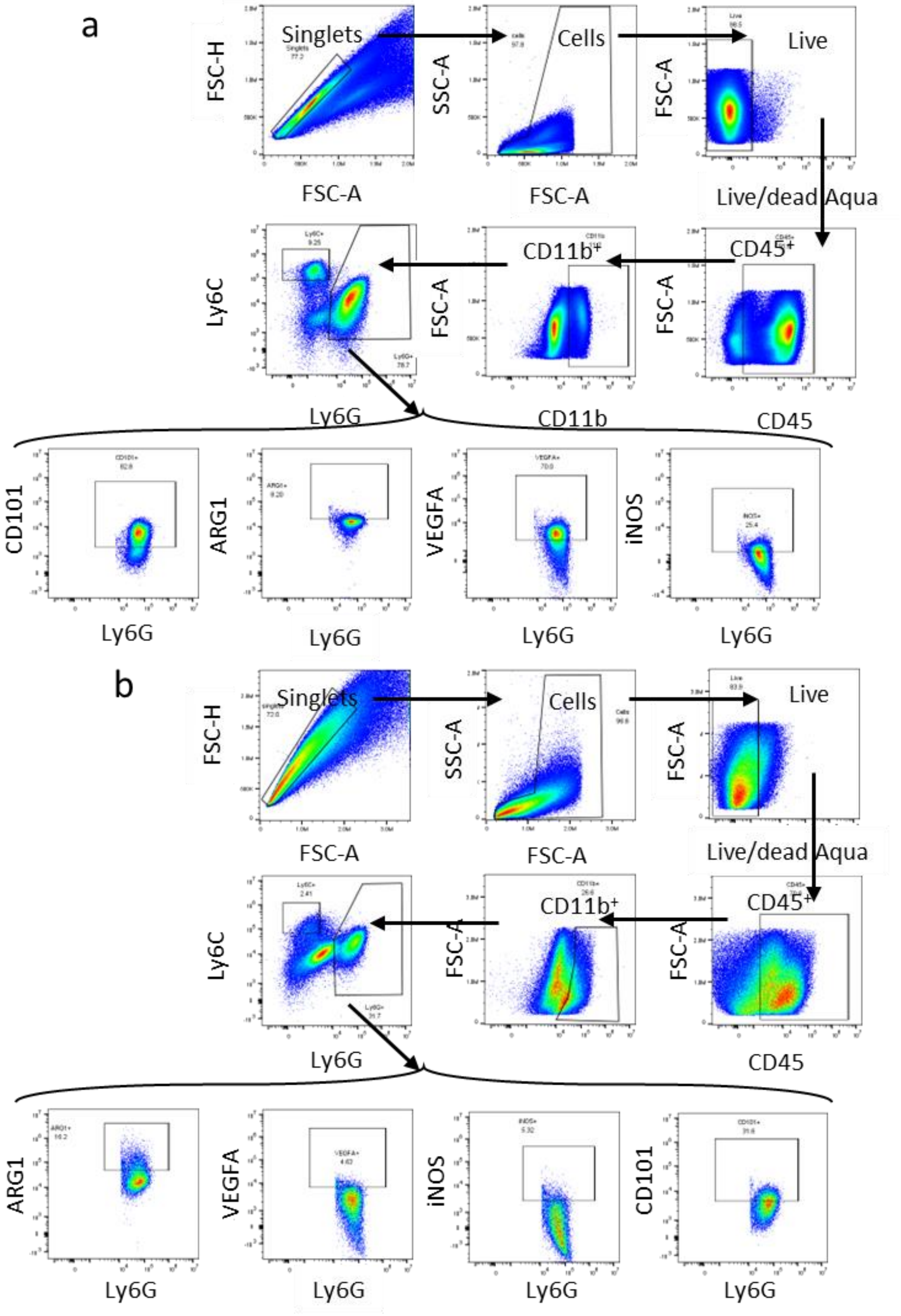


**Supplementary Fig. 12. Gating Strategy for the flow cytometry analysis of myeloid cells.** (**a**) Gating strategy for myeloid cells in the spleen. (**b**) Gating strategy for myeloid cells in the tumor.

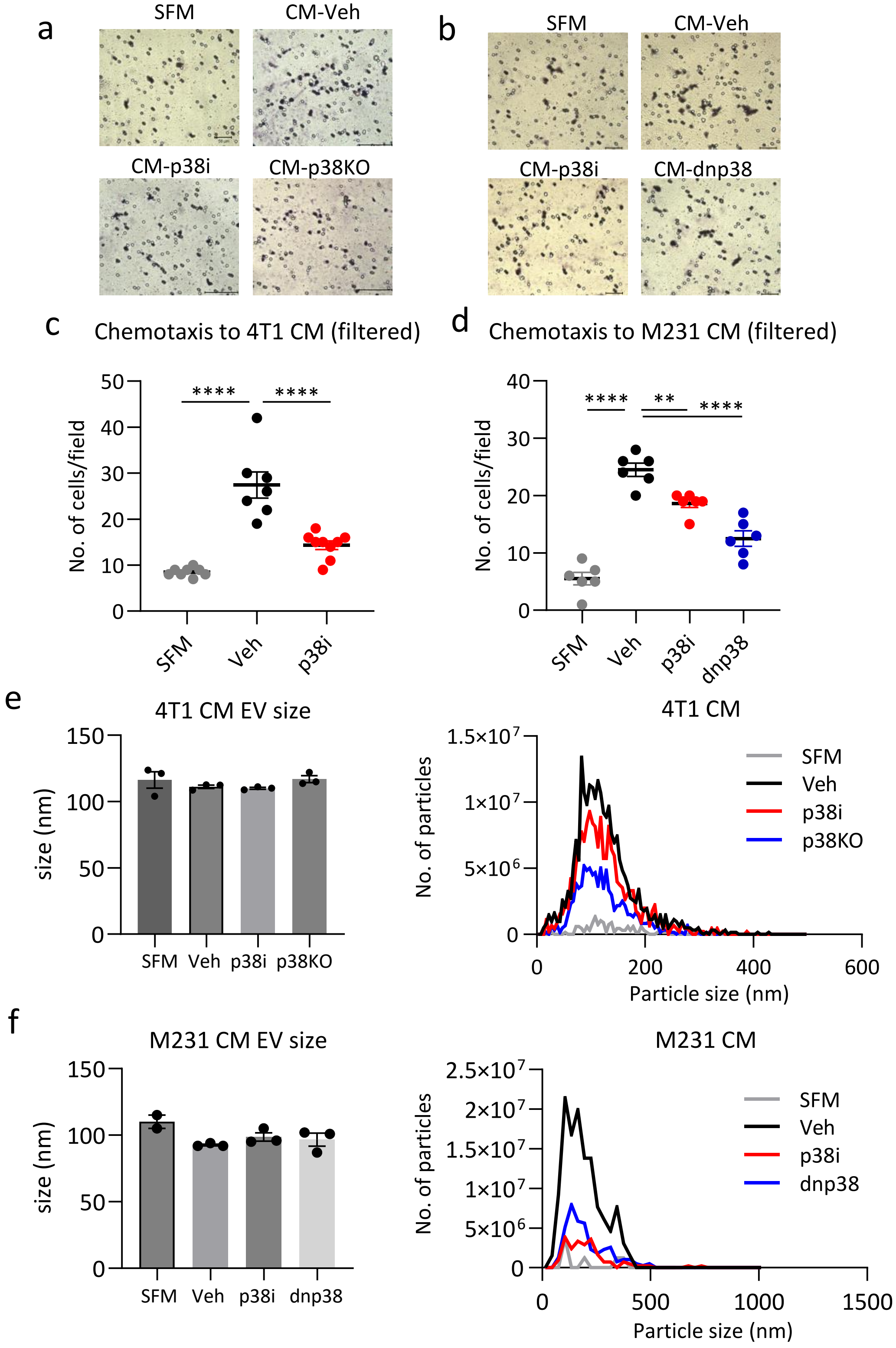


**Supplementary Fig. 13**. Tumor p38 regulates the production of chemotactic cytokines and extra-cellular vesicles (EVs). (**a**) Representative images for chemotaxis to 4T1 CM shown in Fig. 6a. Scale bar represents 50µm. (**b**) Representative images for chemotaxis to MDA-MB-231 CM shown in Fig. 6b. Scale bar represents 50µm. (**c**) Chemotaxis of RAW264.7 cells to 4T1 CM depleted of small molecules using Amicon 3K centrifugal filters. (**d**) Chemotaxis of RAW264.7 cells to MDA-MB-231 CM filtered using Amicon 3K filters. (**e**) Sizes of EVs present in 4T1 CM and size distribution. (**f**) Size of EVs present in MDA-MB-231 CM and size distribution. (Continued on next page).

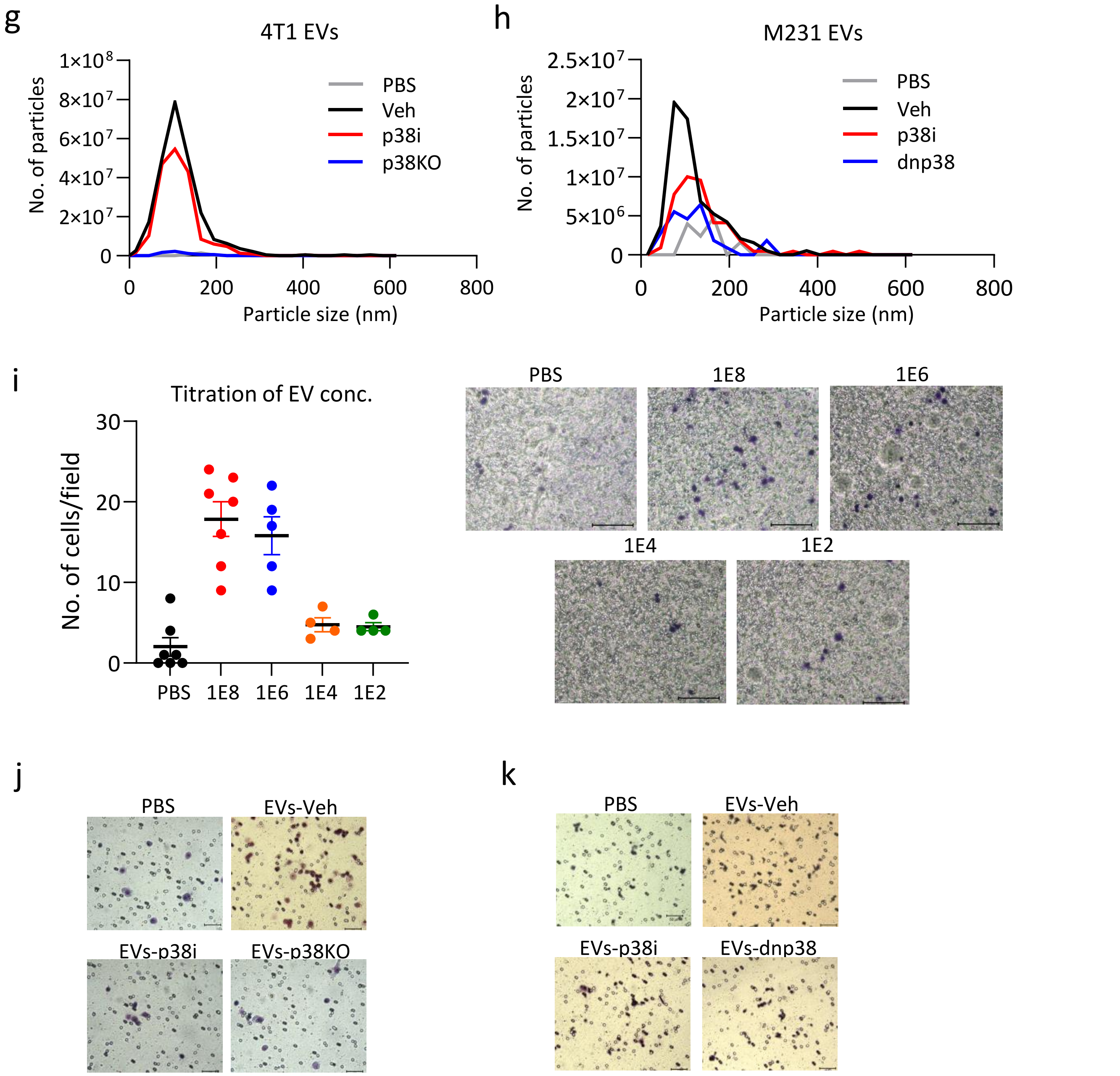


**Supplementary Fig. 13.** (**e**) Size distribution of 4T1 EVs isolated from CM. (**f**) Size distribution of M231 EVs isolated from CM. (**g**) Titration of 4T1 EV concentration for chemotaxis assay. (**h**) Representative images for chemotaxis to 4T1 EVs shown in Fig. 6g. Scale bar represents 50µm. (**i**) Representative images for chemotaxis to M231 EVs shown in Fig. 6h. Scale bar represents 50µm.

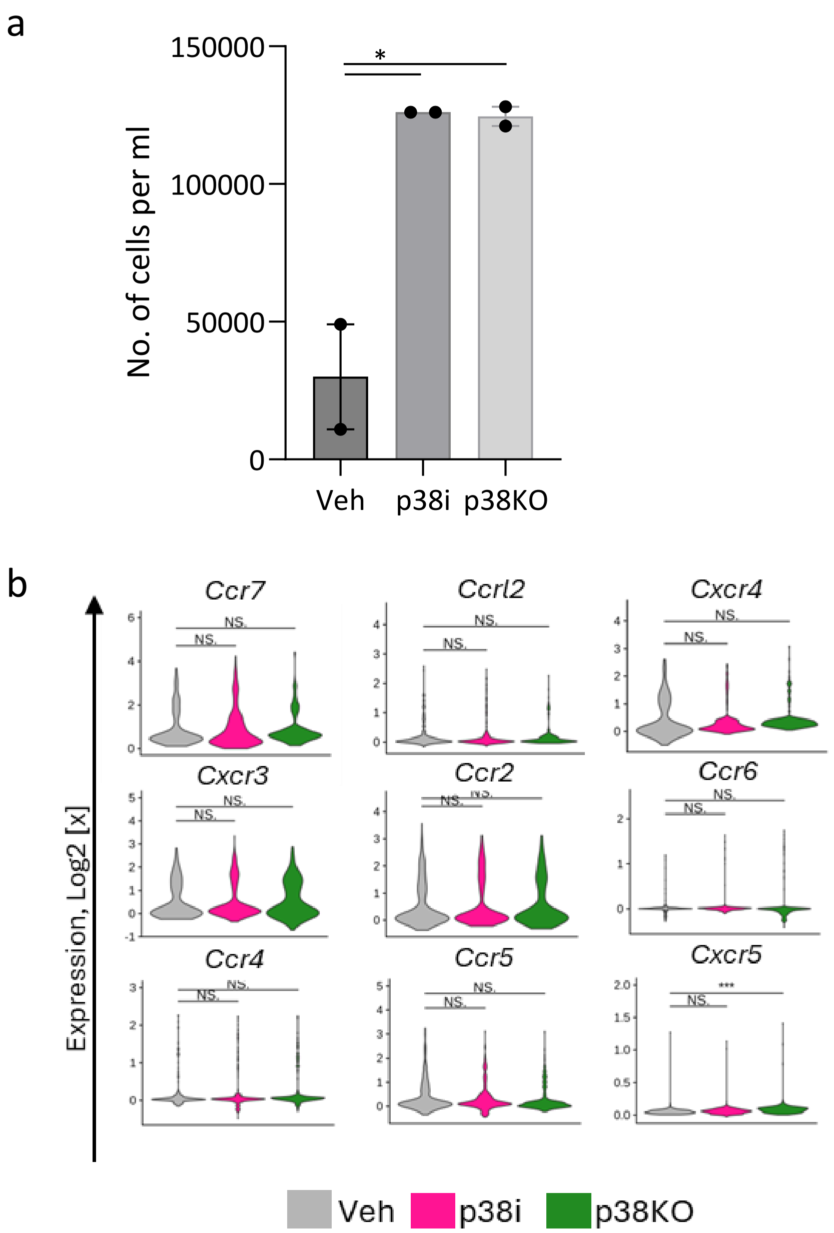


**Supplementary Fig. 14. The analysis of chemotactic migration of T cells to tumor-conditioned media (TCM).** (**a**) Chemotaxis assays with TCM and primary mouse T cells activated by anti-CD3/CD28 magnetic beads and cultured in the presence of IL-2. TCM media (600µl) from vehicle-control, p38i-treated, or p38KO 4T1 cells was added to the bottom chambers 15min prior cell seeding. T cells (~$0.5\times10^{6}$) in 100µl serum-free DMEM were added to the top chamber of each transwell. After 8h, the media from bottom wells was collected and centrifugated. The cell pellets were resuspended in 100µl SFM and cells were counted using automated cell counter. (**b**) Expression of chemokine receptors in T cells was assessed using our scRNAseq data from vehicle-control, p38i treated or p38KO tumors. Violin plots show distribution of mRNA levels (measured in log2 counts per million) of chemokine receptors in the T cells. Comparisons were done by t-test, * $p < 0.05$, ** $p<0.01$, *** $p<0.001$, **** $p<0.0001$, ns = non-significant.

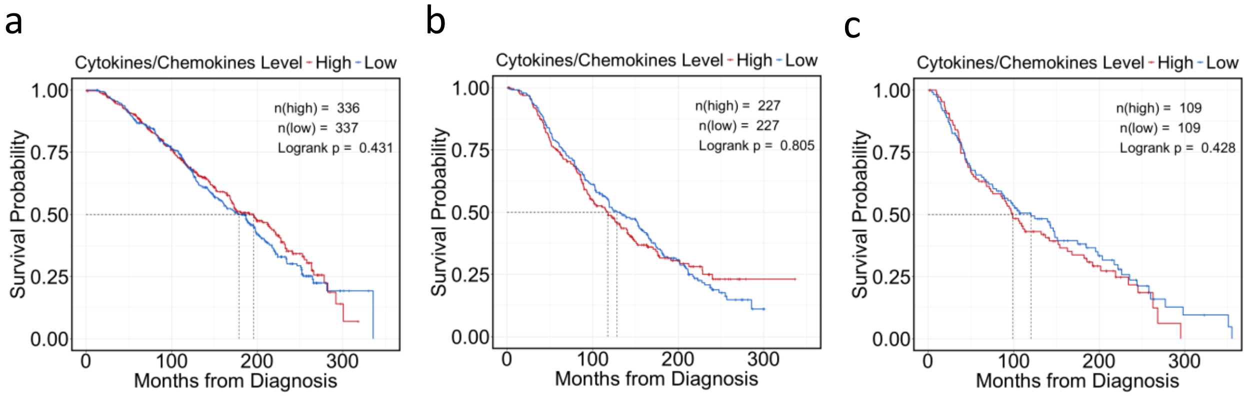


**Supplementary Fig. 15. Survival analysis of breast cancer patients based on expression of p38 regulated factors.** (**a**) Kaplan-Meier survival plot of overall survival of patients with luminal A breast cancer with high (red line) or low (blue line) expression of cytokine-chemokine factors (downregulated by p38i), using the TCGA database. Median survival in the low expression group is 178.56 months vs 195.7 months in the high expression group. (**b**) Kaplan-Meier survival plot of overall survival of patients with luminal B breast cancer with high (red line) or low (blue line) expression of cytokine-chemokine factors (downregulated by p38i), using the TCGA database. Median survival in the low expression group is 138.53 months vs 117.66 months in the high expression group. (**c**) Kaplan-Meier survival plot of overall survival of patients with Her2$^{+}$ breast cancer with high (red line) or low (blue line) expression of cytokine-chemokine factors downregulated by p38i, using the TCGA database. Median survival in the low expression group is 120.13 months vs 98.76 months in the high expression group.
